# Scavenging of reactive dicarbonyls with 2-hydroxybenzylamine reduces atherosclerosis in hypercholesterolemic *Ldlr^−/−^* mice

**DOI:** 10.1101/524884

**Authors:** Huan Tao, Jiansheng Huang, Patricia G. Yancey, Valery Yermalitsky, John L. Blakemore, Youmin Zhang, Lei Ding, Irene Zagol-Ikapitte, Venkataraman Amarnath, Olivier Boutaud, John A. Oates, L. Jackson Roberts, Sean S. Davies, MacRae F. Linton

## Abstract

The pathogenesis of atherosclerosis may be accelerated by oxidative stress, which produces lipid peroxidation. Among the products of lipid peroxidation are highly reactive dicarbonyls including isolevuglandins (IsoLGs) and malondialdehyde (MDA) that covalently modify proteins. We investigated the impact of treatment with the dicarbonyl scavenger, 2-hydroxybenzylamine (2-HOBA) on HDL function and atherosclerosis in hyperlipidemic *Ldlr^−/−^* mice, a model of familial hypercholesterolemia (FH). Compared to mice treated with vehicle, 2-HOBA significantly decreased atherosclerosis in hypercholesterolemic *Ldlr^−/−^* mice by 31% in the proximal aortas and 60% in *en face* aortas, in the absence of changes in blood lipid levels. 2-HOBA reduced MDA content in HDL and LDL. Consuming a western diet increased plasma MDA-apoAI adduct levels in *Ldlr^−/−^* mice. 2-HOBA inhibited MDA-apoAI formation and increased the capacity of the mouse HDL to reduce macrophage cholesterol stores. Importantly, 2-HOBA reduced the MDA- and IsoLG-lysyl content in atherosclerotic aortas in *Ldlr^−/−^* mice. Furthermore, 2-HOBA diminished oxidative stress-induced inflammatory responses in macrophages, reduced the number of TUNEL-positive cells in atherosclerotic lesions by 72%, and decreased serum proinflammatory cytokines. Furthermore, 2-HOBA enhanced efferocytosis and promoted characteristics of stable plaque formation in mice as evidenced by a 69% (p<0.01) reduction in necrotic core and by increased collagen content (2.7-fold) and fibrous cap thickness (2.1-fold). HDL from patients with FH had increased MDA content resulting in a reduced ability of FH-HDL to decrease macrophage cholesterol content versus controls. Our results demonstrate that dicarbonyl scavenging with 2-HOBA has multiple atheroprotective effects on lipoproteins and reduces atherosclerosis in a murine model of FH, supporting its potential as a novel therapeutic approach for the prevention and treatment of human atherosclerotic cardiovascular disease.

**Abbreviations:** 2-HOBA, 2-hydroxybenzylamine; 4-HOBA, 4-hydroxybenzylamine; MDA, malondialdehyde; 4-HNE, 4-hydroxynonenal; IsoLGs, isolevuglandins; HDL, high-density lipoproteins; LDL, lowdensity lipoprotein; LDLR, low-density lipoprotein receptor; ApoAI, apolipoprotein AI; ApoB, apolipoprotein B; ROS, reactive oxygen species; IL, interleukin.

## Introduction

Atherosclerosis, the underlying cause of heart attack and stroke, is the most common cause of death and disability in the industrial world ^1^. Elevated levels of apolipoprotein B (LDL and VLDL) containing lipoproteins and low levels of HDL increase the risk of atherosclerosis ^1^. Although lowering LDL with HMG-CoA reductase inhibitors has been shown to reduce the risk of heart attack and stroke in large outcomes trials, substantial residual risk for cardiovascular events remains ^2^. Atherosclerosis is a chronic inflammatory disease with oxidative stress playing a critical role ^3, 4^. Oxidative modification of apoB containing lipoproteins enhances internalization leading to foam cell formation ^1, 5^. In addition, oxidized LDL induces inflammation, immune cell activation, and cellular toxicity ^1, 5^. HDL protects against atherosclerosis via multiple roles including promoting cholesterol efflux, preventing LDL oxidation, maintaining endothelial barrier function, and by minimizing cellular oxidative stress and inflammation ^1, 4,6^. HDL-C concentration is inversely associated with cardiovascular disease (CVD) ^6^, but recent studies suggest that assays of HDL function may provide new independent markers for CVD risk ^7, 8^. Evidence has mounted that oxidative modification of HDL compromises its functions, and studies suggest that oxidized HDL is indeed proatherogenic ^1, 6, 9^.

During lipid peroxidation, highly reactive dicarbonyls, including 4-oxo-nonenal (4-ONE) malondialdehyde (MDA) and isolevuglandins (IsoLGs) are formed. These reactive lipid dicarbonyls covalently bind to DNA, proteins, and phospholipid causing alterations in lipoprotein and cellular functions ^1, 10, 11^. In particular, modification with reactive lipid dicarbonyls promotes inflammatory responses and toxicity that may be relevant to atherosclerosis ^12, 13, 14, 15^. Identifying effective strategies to assess the contribution of reactive lipid dicarbonyls to disease processes in vivo has been challenging. Although formation of reactive lipid species, including dicarbonyls, theoretically could be suppressed simply by lowering levels of reactive oxygen species (ROS) using dietary antioxidants, the use of antioxidants to prevent atherosclerotic cardiovascular events has proven problematic with most clinical outcomes trials failing to show a benefit ^1, 16^. Dietary antioxidants like vitamin C and vitamin E are relatively ineffective suppressors of oxidative injury and lipid peroxidation. In fact, careful studies of patients with hypercholesterolemia found that the doses of vitamin E required to significantly reduce lipid peroxidation were substantially greater than those typically used in most clinical trials ^17^. Furthermore, the high doses of antioxidants needed to suppress lipid peroxidation have been associated with significant adverse effects, likely because ROS play critical roles in normal physiology, including protection against bacterial infection and in a number of cell signaling pathways. Finally, for discovery purposes, the use of antioxidants provides little information about the role of reactive lipid dicarbonyls because suppression of ROS inhibits formation of a broad spectrum of oxidatively modified macromolecules besides reactive lipid dicarbonyl species.

An alternative approach to broad suppression of ROS utilizing antioxidants is to use small molecule scavengers that selectively react with lipid dicarbonyl species without altering ROS levels, thereby preventing reactive lipid dicarbonyls from modifying cellular macromolecules without disrupting normal ROS signaling and function. 2-hydroxybenzylamine (2-HOBA) rapidly reacts with lipid dicarbonyls such as IsoLG, ONE, and MDA, but not with lipid monocarbonyls such as 4-hydroxynonenal ^15, 18, 19, 20^. The 2-HOBA isomer 4-hydroxybenzylamine (4-HOBA) is ineffective as a dicarbonyl scavenger ^21^. Both of these compounds are orally bioavailable, so they can be used to examine the effects of lipid dicarbonyl scavenging in vivo ^13, 22^. 2-HOBA protects against oxidative stress associated hypertension ^13^, oxidant induced cytotoxicity ^15^, neurodegeneration, ^14^ and rapid pacing induced amyloid oligomer formation ^23^. While there is evidence that reactive lipid dicarbonyls play a role in atherogenesis ^6, 7^, to date the effects of scavenging lipid dicarbonyl on the development of atherosclerosis have not been examined.

Here, for the first time, we show that 2-HOBA treatment significantly attenuates atherosclerosis development in hypercholesterolemic *Ldlr^−/−^* mice. Importantly, 2-HOBA treatment inhibits cell death and necrotic core formation in lesions, leading to the formation of characteristics of more stable plaques as evidenced by increased lesion collagen content and fibrous cap thickness. Consistent with the decrease in atherosclerosis from 2-HOBA treatment being due to scavenging of reactive dicarbonyls, the atherosclerotic lesion MDA and IsoLG adduct content was markedly reduced in 2-HOBA treated versus control mice. We further show that 2-HOBA treatment results in decreased MDA-LDL and MDA-HDL. In addition, MDA-apoAI adduct formation was decreased, and importantly, 2-HOBA treatment caused more efficient HDL function in reducing macrophage cholesterol stores. Thus, scavenging of reactive carbonyls with 2-HOBA has multiple antiatherogenic therapeutic effects that likely contribute to its ability to reduce the development of atherosclerosis in hypercholesterolemic *Ldlr^−/−^* mice. We also found that HDL from humans with severe familial hypercholesterolemia (FH) contained increased MDA adducts versus control subjects, and that FH-HDL were extremely impaired in reducing macrophage cholesterol stores. Taken together, our studies raise the possibility that reactive dicarbonyl scavenging is a novel therapeutic approach to prevent and treat human atherosclerosis.

## Results

### 2-HOBA treatment attenuates atherosclerosis without altering plasma cholesterol levels in *Ldlr^−/−^* mice

Eight week old, female *Ldlr^−/−^* mice were fed a western-type diet for 16 weeks and were continuously treated with vehicle alone (water) or water containing either 2-HOBA or 4-HOBA, a nonreactive analogue. Treatment with 2-HOBA reduced the extent of proximal aortic atherosclerosis by 31.1% and 31.5%, compared to treatment with either vehicle or 4-HOBA, respectively (Figures 1A and 1B). In addition, en face analysis of the aorta demonstrated that treatment of female *Ldlr^−/−^* mice with 2-HOBA reduced the extent of aortic atherosclerosis by 60.3% and 59.1% compared to administration of vehicle and 4-HOBA, respectively (Figures 1C and 1D). Compared to administration of vehicle or 4-HOBA, 2-HOBA treatment did not affect body weight, water consumption, or diet uptake (Supplemental Figures 1A-1C). In addition, the plasma total cholesterol and triglyceride levels were not significantly different (Figure 1E), and the lipoprotein distribution was similar between the 3 groups of mice (Supplemental Figure 1D). Consistent with these results, treatment of male *Ldlr^−/−^* mice with 1g/L of 2-HOBA, under similar conditions of being fed a western diet for 16 weeks, reduced the extent of proximal aortic and whole aorta atherosclerosis by 37% and 45%, respectively, compared to treatment with water (Supplemental Figure 2A-2D) but did not affect the plasma total cholesterol levels (Supplemental Figure 2E). A similar reduction in atherosclerosis was observed when male *Ldlr^−/−^* mice were treated with 3 g/L of 2-HOBA (Data not shown). Thus, for the first time, we demonstrate that 2-HOBA treatment significantly decreases atherosclerosis development in an experimental mouse model of FH without changing plasma cholesterol and triglyceride levels. Examination of the proximal aortic MDA adduct content by immunofluorescence staining using an antibody against MDA-protein adducts (Abcam cat# ab6463) shows that the MDA adduct levels were reduced by 68.5% and 66.8% in 2-HOBA treated mice compared to mice treated with vehicle alone or 4-HOBA (Figures 2A and 2B). We determined that the anti-MDA-protein antibody does not recognize either free MDA or MDA-2-HOBA adducts (Supplemental Figures 3A and 3B). In addition, 2-HOBA does not interfere with the antibody recognition of MDA-albumin adducts (Supplemental Figures 3A and 3B). Quantitative measurement of the whole aorta MDA- and IsoLG-lysyl adducts by LC/MS/MS demonstrates that compared to 4-HOBA treatment, administration of 2-HOBA decreased the MDA and IsoLG adduct content by 59% and 23%, respectively (Figures 2C and 2D). We determined by LC/MS/MS that the plasma levels of 2-HOBA in the male *Ldlr^−/−^* mice after 16 weeks of treatment with 1g of 2-HOBA/L of water were 469 ± 38 ng/mL, which is similar to what we previously reported in C57BL6 mice receiving 1g/L of 2-HOBA^22^. In addition, these levels are in the same range as the plasma 2-HOBA levels in humans in a recent safety trial ^24^. We determined that 4-HOBA was cleared more readily as compared to 2-HOBA. The plasma levels of 2-HOBA versus 4-HOBA in male *Ldlr^−/−^* mice 30min after oral gavage of 5mg were 3.6-fold higher (Supplemental Figure 4A). A similar difference in plasma levels was observed in C57BL6 mice 30min after intraperitoneal injection with 2-HOBA versus 4-HOBA (Supplemental Figure 4B). Importantly, IsoLG-2-HOBA adducts (with masses consistent with potential keto-pyrrole, anhydro-lactam, keto-lactam, pyrrole, and anhydro-hydroxylactam adducts) were present in the hearts and livers of *Ldlr^−/−^* mice after 16 weeks on a western diet, whereas IsoLG-4-HOBA adducts were nondetectable (Supplemental Figure 5 and Table). In addition, the MDA-2-HOBA versus MDA-4-HOBA adducts (with mass consistent with propenal-HOBA adducts) were markedly increased in the urine collected during 18h after oral gavage (5mg) treatment of *Ldlr^−/−^* mice after 16 weeks on a western diet (Supplemental Figures 6A-6B). The liver, kidney, and spleen from 2-HOBA versus 4-HOBA treated *Ldlr^−/−^* mice also contained 3-, 5-, and 11-fold more propenal-HOBA adducts (Supplemental Figures 6C-6E). These results are consistent with the 2-HOBA effects on atherosclerosis mainly being due to reactive lipid dicarbonyl scavenging. Indeed, the effects were not due to general inhibition of lipid peroxidation or metal ion chelating as the urine F_2_-isoprostane levels were not different in *Ldlr^−/−^* mice treated with vehicle, 4-HOBA, and 2-HOBA (Supplemental Figure 7).

**Figure 1.**
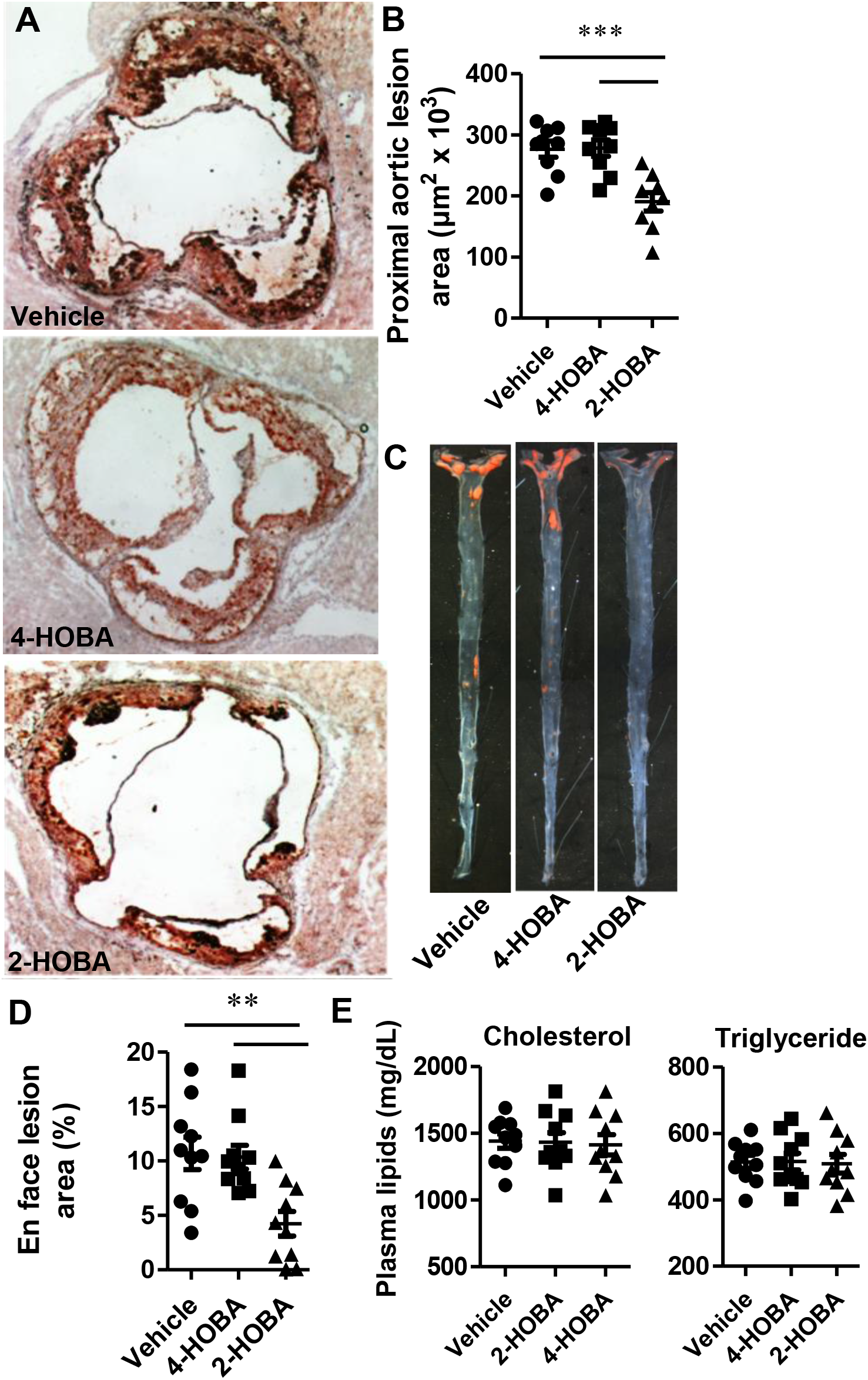
2-HOBA attenuates atherosclerosis in hypercholesterolemic female *Ldlr^−/−^* mice. 8-week old *Ldlr^−/−^* female mice were pretreated with 1 g/L 2-HOBA or 1 g/L 4-HOBA (nonreactive analogue) or vehicle (water) for 2 weeks and then the treatment was continued for 16 weeks during which the mice were fed a Western diet. (A and C) Representative images show Red-Oil-O stain in proximal aorta root sections (A) and in open-pinned aortas (C). (B and D) Quantitation of the mean Oil-Red-O stainable lesion area in aorta root sections (B) and en face aortas (D). N = 9 or 10 per group, ** p<0.01, *** p<0.001. (E) The plasma total cholesterol and triglyceride levels. N = 9 or 10 per group. The *p* value was analyzed by One-way ANOVA with Bonferroni’s post-test.

**Figure 2.**
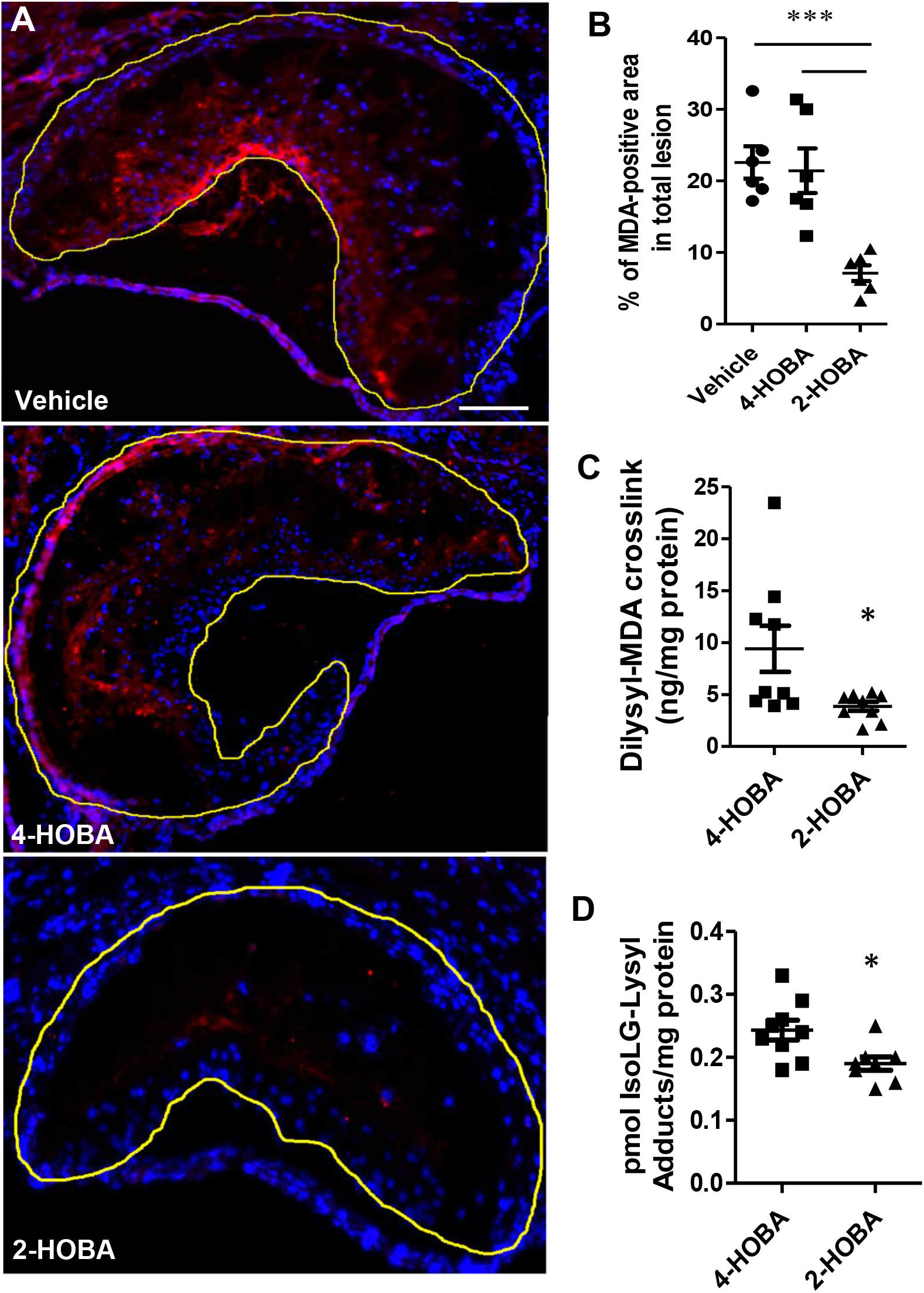
2-HOBA decreases the MDA adduct content of proximal aortic atherosclerotic lesions in *Ldlr^−/−^* mice. MDA was detected by immunofluorescence using anti-MDA primary antibody and fluorescent-labeled secondary antibody. Nuclei were counterstained with Hoechst (Blue). (A) Representative images show MDA staining (Red) in proximal aortic root sections. (B) Quantitation of the mean MDA positive lesion area in aortic root sections using ImageJ software. Data are expressed as mean ± SEM, N = 6 per group, *** p<0.001. One-way ANOVA with Bonferroni’s post-test. (C-D). Aortic tissues were isolated from the *Ldlr^−/−^* mice and Dilysyl-MDA crosslinks (C) or IsoLG-Lysyl (D) were measured by LC/MS/MS. Data are presented as mean ± SEM, N = 8 or 9 per group, * p<0.05, *p* values were analyzed by Mann-Whitney test.

### 2-HOBA treatment promotes formation of characteristics of more stable atherosclerotic plaques in hypercholesterolemic *Ldlr^−/−^* mice

As vulnerable plaques exhibit higher risk for acute cardiovascular events in humans ^1^, we examined the effects of 2-HOBA treatment on characteristics of plaque stabilization by quantitating the atherosclerotic lesion collagen content, fibrous cap thickness, and necrotic cores (Figures 3A-3D). Compared to administration of vehicle or 4-HOBA, 2-HOBA treatment increased the collagen content of the proximal aorta by 2.7- and 2.6-fold respectively (Figures 3A and 3B). In addition, the fibrous cap thickness was 2.31– and 2.29–fold greater in lesions of 2-HOBA treated mice versus vehicle and 4-HOBA treated mice (Figures 3A and 3C). Importantly, the % of necrotic area in the proximal aorta was decreased by 74.8% and 73.5% in mice treated with 2-HOBA versus vehicle and 4-HOBA (Figures 3A and 3D). Taken together, these data show that 2-HOBA suppresses the characteristics of vulnerable plaque formation in the hypercholesterolemic *Ldlr^−/−^* mice.

**Figure 3.**
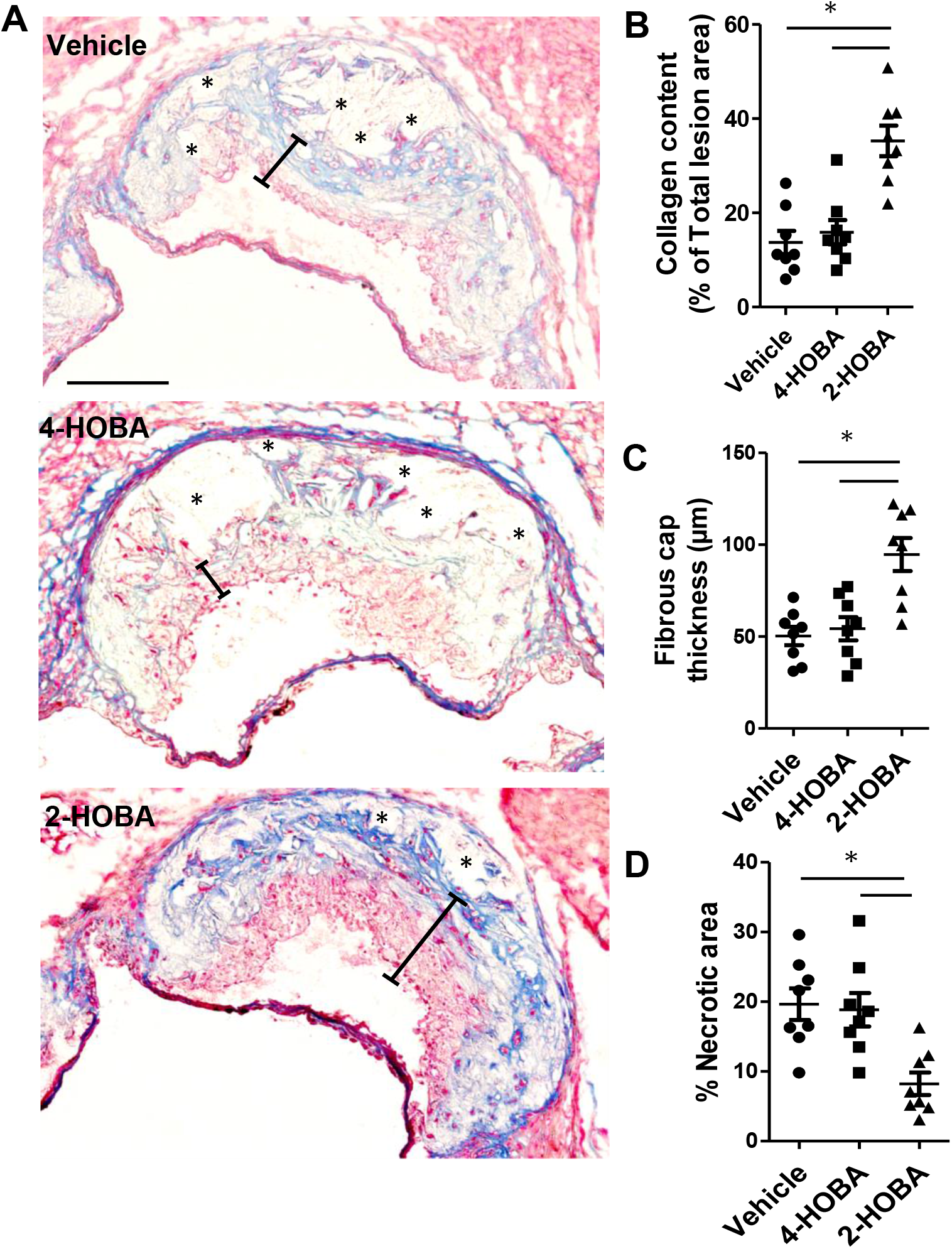
2-HOBA promotes features of stable atherosclerotic plaques in *Ldlr^−/−^* mice. Masson’s Trichrome stain was done to analyze characteristics associated with atherosclerotic lesion stability in proximal aorta sections of *Ldlr^−/−^* mice. (A) Representative images show Masson’s Trichrome stain in aorta root sections. The collagen content (B), fibrous cap thickness (C), and necrotic area (D) were quantitated using ImageJ software. N = 8 per group. * p< 0.05, One-way ANOVA with Bonferroni’s post-test. Scale bar = 100 μm. Blue shows collagen, Red, cytoplasm, Black, nuclei.

### 2-HOBA treatment promotes cell survival and efferocytosis and reduces inflammation

As enhanced cell death and insufficient efferocytosis promote necrotic core formation and destabilization of atherosclerotic plaques, we next examined the effects of 2-HOBA treatment on cell death and efferocytosis in atherosclerotic lesions in the proximal aorta (Figures 4A-4D). Compared to treatment with either vehicle or 4-HOBA, the number of TUNEL positive cells was reduced by 72.9% and 72.4% in the proximal aortic lesions of 2-HOBA treated mice (Figures 4A and 4C). Consistent with reactive lipid dicarbonyl scavenging maintaining efficient efferocytosis, the number of TUNEL positive cells not associated with macrophages was increased by 1.9- and 2.0-fold in lesions of mice treated with vehicle and 4-HOBA versus 2-HOBA (Figures 4B and 4D). Consistent with lesion necrosis being linked to enhanced inflammation, the serum levels of IL-1β, IL-6, TNF-α, and serum amyloid A were reduced in 2-HOBA versus 4-HOBA or vehicle treated *Ldlr^−/−^* mice (Figure 5), suggesting that reactive dicarbonyl scavenging decreased systemic inflammation. As studies have demonstrated that incubation of cells with either H_2_O_2_ ^15, 25, 26^ or oxidized LDL^27, 28, 29^ induces lipid peroxidation, inflammation, and death, we next determined the in vitro effects of reactive dicarbonyl scavenging on the cellular response to oxidative stress. Examination of the susceptibility of macrophages and endothelial cells to apoptosis in response to H_2_O_2_ treatment demonstrates that compared to incubation with vehicle or 4-HOBA, 2-HOBA markedly decreased the number of apoptotic cells in both macrophage and endothelial cell cultures (Figures 6A and 6B). In addition, 2-HOBA treatment significantly reduced the macrophage inflammatory response to oxidized LDL as shown by the decreased mRNA levels of IL-1β, IL-6, and TNF-α (Figures 6C-6E). Similar results were observed for the impact of 2-HOBA on the inflammatory cytokine response of macrophages treated with H_2_O_2_ versus vehicle or 4-HOBA treatment (Figures 6F-6H). Consistent with the 2-HOBA effects on cell death and inflammation being due to scavenging reactive dicarbonyls, the levels of MDA-2-HOBA versus MDA-4-HOBA adducts were increased in medium from cells treated with oxidized LDL (Supplemental Figure 8). In addition, 2-HOBA did not have a direct effect on prosurvival, antiinflammatory signaling in the absence of oxidative stress,^30^ as there was no difference in pAKT levels in macrophages treated with vehicle, 4-HOBA, and 2-HOBA in the absence and presence of insulin (Supplemental Figure 9). Due to the striking impact on inflammatory cytokines in vivo, we also measured urinary prostaglandins to evaluate whether 2-HOBA might be inhibiting cyclooxygenase (COX). Urine samples were analyzed for 2,3-dinor-6-keto-PGF1, 11-dehydro TxB2, PGE-M, PGD-M by LC/MS. We found that there were no significant differences in levels of these major urinary prostaglandin metabolites of *Ldlr^−/−^* mice treated with 2-HOBA compared to the vehicle control (Supplemental Figure 10), indicating that 2-HOBA was not significantly inhibiting COX in vivo in mice. Taken together, these data show that 2-HOBA treatment maintains efficient efferocytosis in vivo and prevents apoptosis and inflammation in response to oxidative stress by scavenging reactive dicarbonyls.

**Figure 4.**
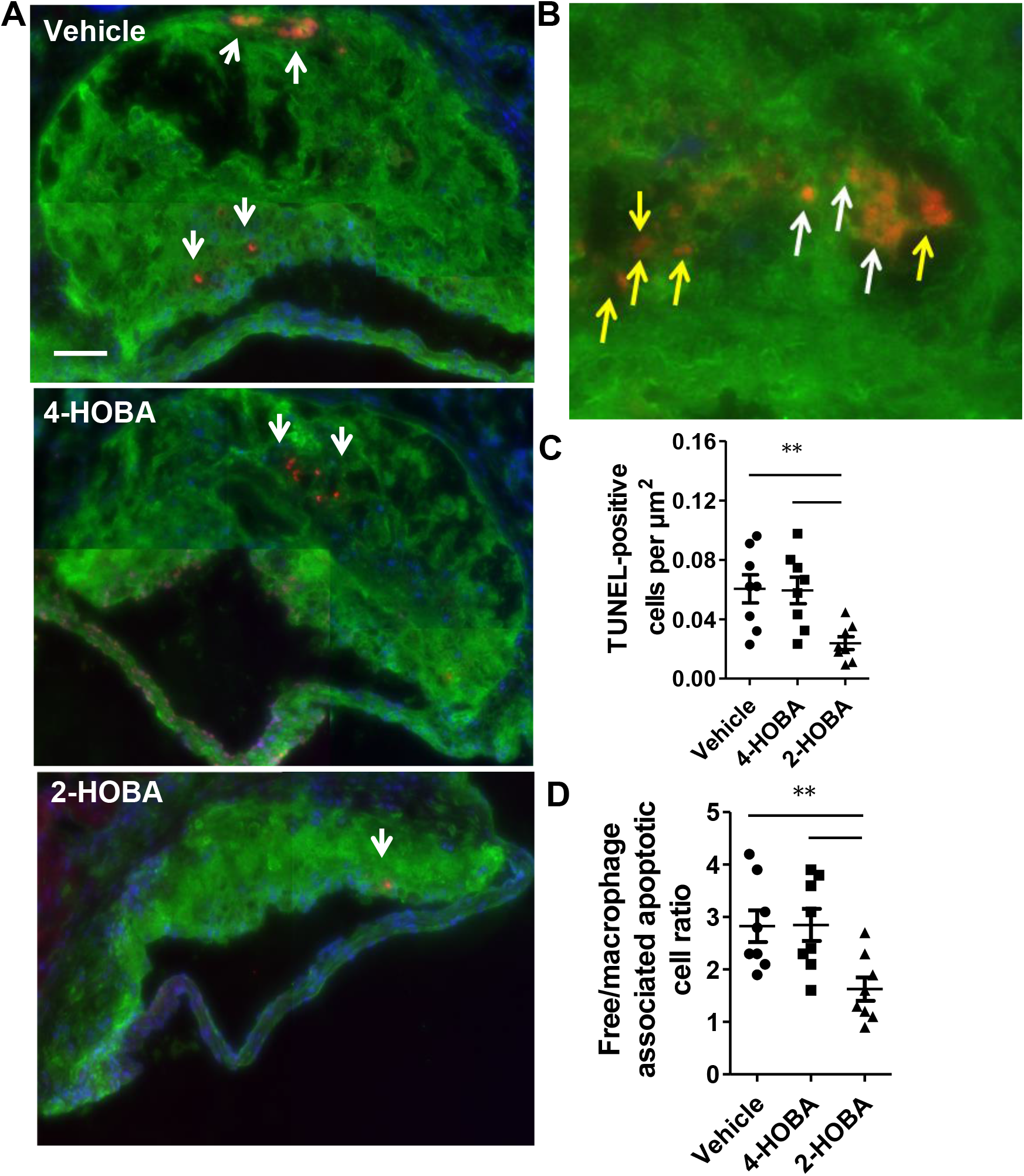
2-HOBA prevents cell death and increases efferocytosis in atherosclerotic lesions of *Ldlr^−/−^* mice. (A) Representative images show dead cells that were detected by TUNEL staining (Red) of proximal aorta sections. Macrophages were detected by anti-macrophage primary antibody (green), and nuclei were counterstained with Hoechst (blue). (B) A representative image taken at a higher magnification to indicate macrophage-associated TUNEL stain (yellow arrows) and white arrows indicate free dead cells that were not associated with macrophages. (C) Quantitation of the number of TUNEL-positive nuclei in proximal aortic sections. (D) Efferocytosis was examined by quantitating the free versus macrophage-associated TUNEL-positive cells in the proximal aortic sections. Data are expressed as mean ± SEM (N = 8 per group). Scale bar = 50 μm, ** p<0.01, One-way ANOVA with Bonferroni’s post-test.

**Figure 5.**
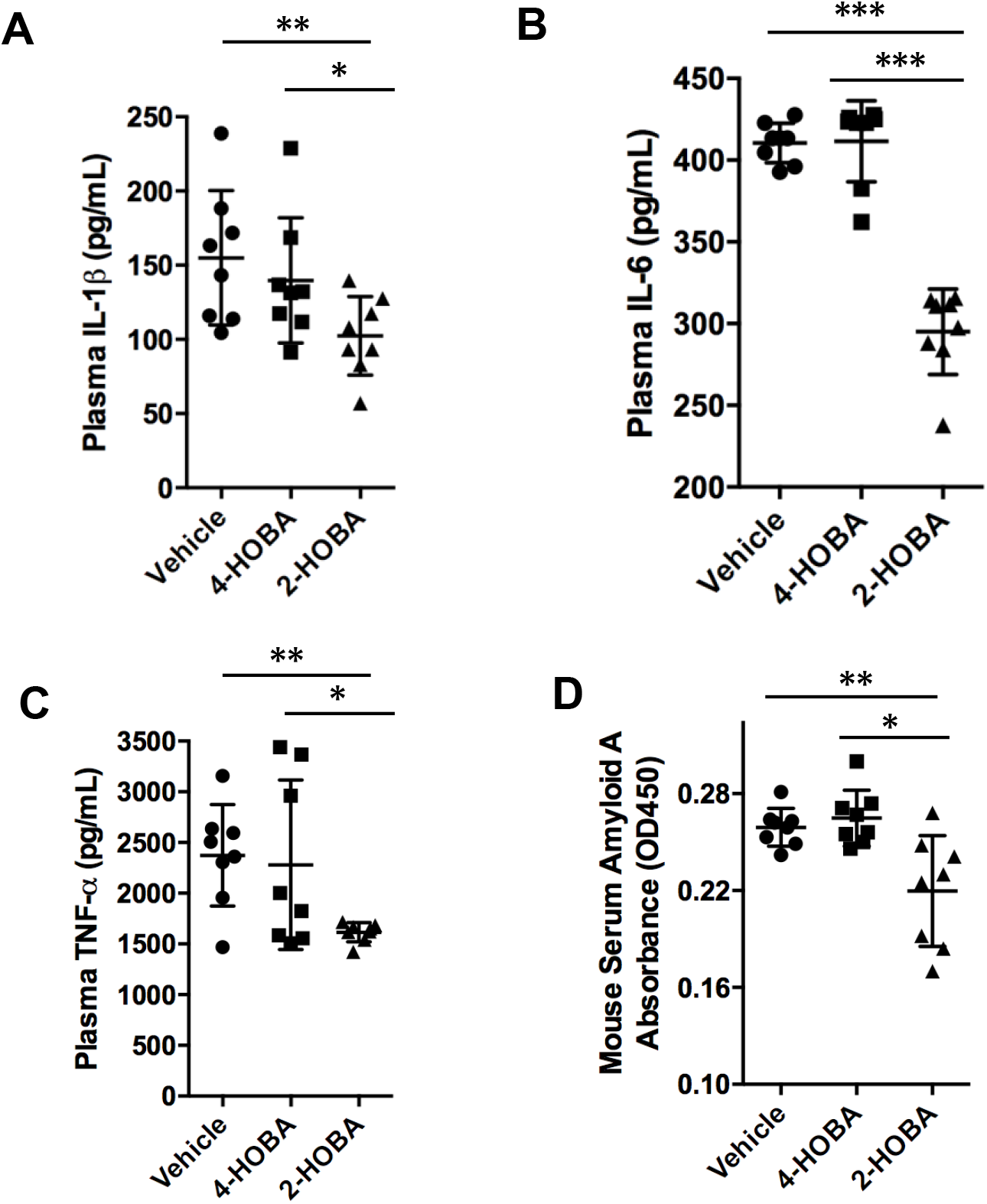
2-HOBA reduces the plasma inflammatory cytokines in hypercholesterolemic *Ldlr^−/−^* mice. The inflammatory cytokines including IL-1β (A), IL-6 (B), TNF-α (C) and SAA (D) were measured by ELISA in plasma from mice consuming a western diet for 16 weeks and treated with 2-HOBA, 4-HOBA, or vehicle. N = 8 per group. *p<0.05, **p<0.01. *** p<0.001, One-way ANOVA with Bonferroni’s post-test.

**Figure 6.**
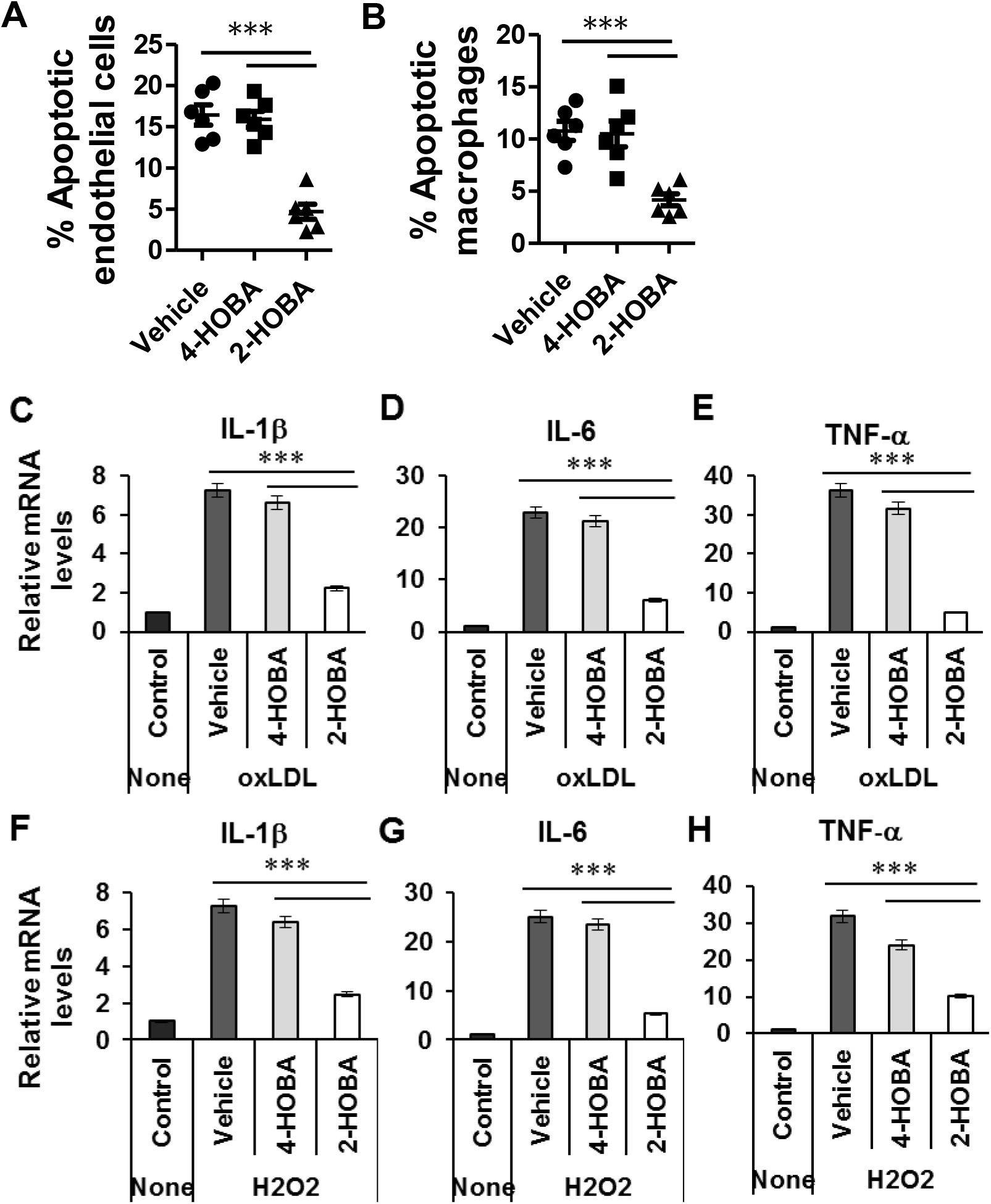
In vitro treatment with 2-HOBA suppresses oxidative stress-induced cell apoptosis and inflammation. (A and B) Mouse aortic endothelial cells (A) or primary macrophages (B) were incubated for 24h with 250 μM H_2_O_2_ alone or with either 4-HOBA or 2-HOBA (500 μM). Apoptotic cells were then detected by Annexin V staining and flow cytometry. (C to H) The mRNA levels of IL-1β, IL-6, and TNF-α were analyzed by real time PCR in the peritoneal macrophages incubated for 24 h with either oxidized LDL (C-E) or 250 μM H_2_O_2_ (F-H) alone or with either 4-HOBA or 2-HOBA (500 μM). (A to H) Data are presented as mean ± SEM from three independent experiments, *** p<0.001, One-way ANOVA with Bonferroni’s post-test.

### Effects of 2-HOBA on MDA modification and function of lipoproteins and the impact of familial hypercholesterolemia on lipoprotein MDA adduct content and function

Treatment of the *Ldlr^−/−^* mice fed a western diet for 16 weeks with 2-HOBA versus 4-HOBA or vehicle decreased the plasma levels of MDA (Supplemental Figure 11A). Compared to treatment with either vehicle or 4-HOBA, the MDA adduct content in isolated LDL measured by ELISA was reduced by 57% and 54%, respectively, in *Ldlr^−/−^* mice treated with 2-HOBA (Supplemental Figure 11B). By comparison, LDL from control and FH subjects contained similar amounts of MDA adducts, which were not significantly different (Supplemental Figure 11C). MDA modification of LDL induces foam cell formation and examination of the ability of LDL from 2-HOBA versus 4-HOBA or vehicle treated *Ldlr^−/−^* mice to enrich cells with cholesterol was not different (Supplemental Figure 11D). Similar results were observed with FH versus control LDL (Data not shown). This observation was due to the plasma LDL from FH subjects and hypercholesterolemic *Ldlr^−/−^* mice being insufficiently modified with MDA to induce cholesterol loading as we determined by in vitro modification of LDL that the MDA content must be 2500 ng/mg LDL protein to enrich cells with cholesterol. As oxidative modification of HDL impairs its functions, we next examined the effects of 2-HOBA treatment on HDL MDA content and function. Treatment of *Ldlr^−/−^* mice with 2-HOBA reduced the MDA adduct content of isolated HDL as measured by ELISA by 57% and 56% (Figure 7A) compared to treatment with either vehicle or 4-HOBA. Next, we examined the effects on apoAI MDA adduct formation. ApoAI was isolated from plasma by immunoprecipitation, and MDA-apoAI was detected by western blotting with the antibody to MDA-protein adducts. After 16 weeks on the western-type diet, *Ldlr^−/−^* mice treated with vehicle or 4-HOBA had markedly increased plasma levels of MDA-apoAI compared to *Ldlr^−/−^* mice consuming a chow diet (Figures 7B and 7C). In contrast, treatment of *Ldlr^−/−^* mice consuming a western diet with 2-HOBA dramatically reduced plasma MDA-apoAI adducts (Figures 7B and 7C). The levels of apoAI were similar among the 4 groups of mice (Figure 7B). Importantly, the HDL isolated from 2-HOBA treated *Ldlr^−/−^* mice was 2.2- and 1.7-fold more efficient at reducing cholesterol stores in *Apoe^−/−^* macrophage foam cells versus vehicle and 4-HOBA treated mice (Figure 7D). In addition, HDL from human subjects with severe FH pre- and post-LDL apheresis (LA) had 5.9-fold and 5.6-fold more MDA adducts compared to control HDL as measured by ELISA (Figure 7E). We also found that the dilysyl-MDA crosslink levels as measured by LC/MS/MS were higher in HDL from FH versus control subjects (Figure 7F). Importantly, HDL from FH versus control subjects lacked the ability to reduce the cholesterol content of cholesterol-enriched *Apoe^−/−^* macrophages (Figure 7G). While the effects of MDA modification of lipid-free apoAI on cholesterol efflux are established,^31^ studies are controversial regarding the impact of modification of HDL^32, 33^. Therefore, we determined the impact of in vitro modification of HDL with MDA on the the ability of HDL to reduce the cholesterol content of macrophage foam cells as it relates to the MDA adduct content measured by ELISA (Supplemental Figure 12A and 12B). MDA modification of HDL inhibited the net cholesterol efflux capacity in a dose dependent manner, and importantly the MDA-HDL adduct levels which impacted the cholesterol efflux function were in the same range as MDA adduct levels in HDL from FH subjects and hypercholesterolemic *Ldlr^−/−^* mice. Taken together, dicarbonyl scavenging with 2-HOBA prevents macrophage foam cell formation by improving HDL net cholesterol efflux capacity. In addition, our studies suggest that scavenging of reactive lipid dicarbonyls could be a relevant therapeutic approach in humans given that HDL from subjects with homozygous FH contain increased MDA and IsoLG and enhanced foam cell formation.

**Figure 7.**
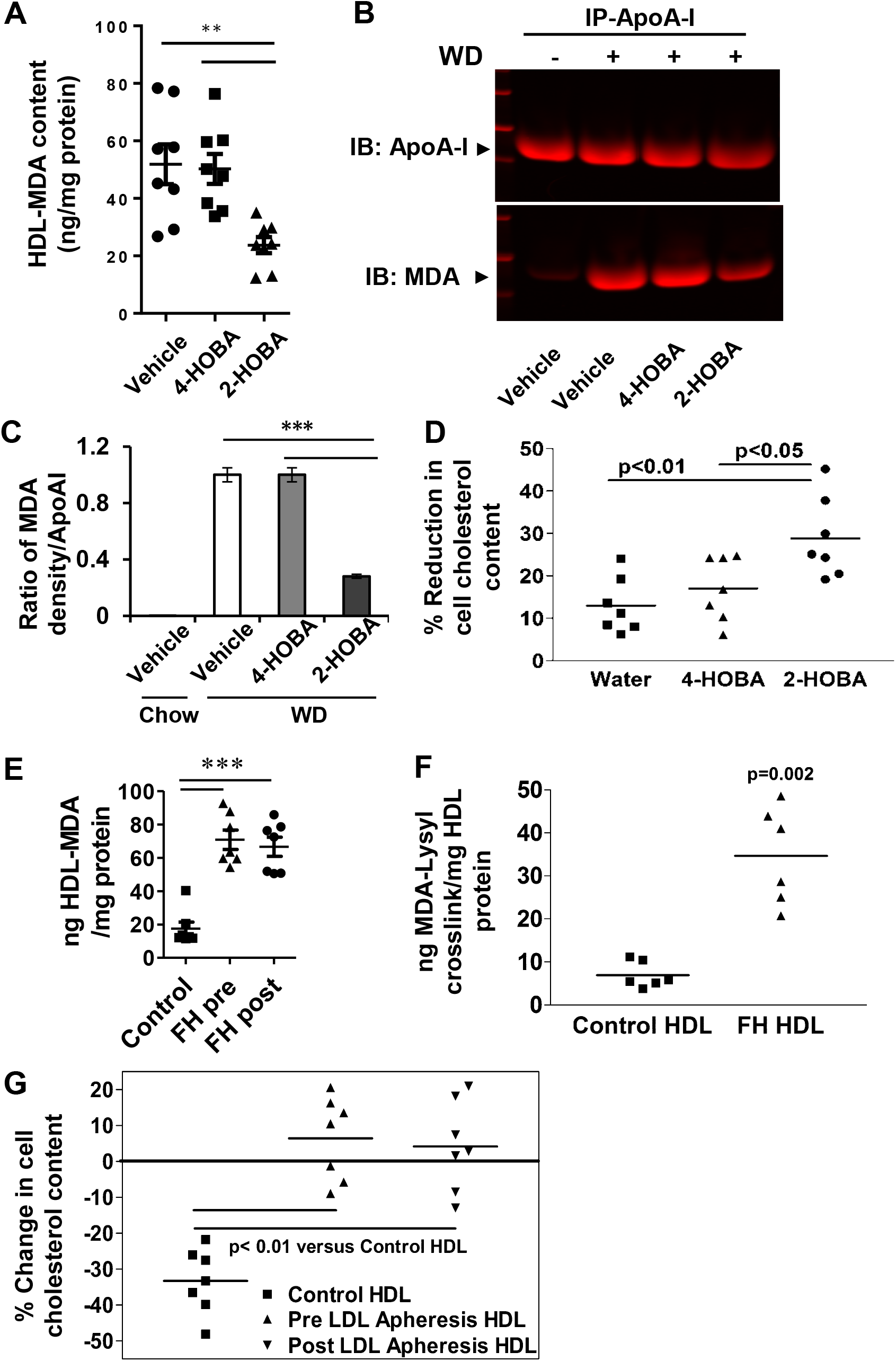
Effects of 2-HOBA on MDA-HDL adducts and HDL function. (A) The levels of MDA adducts were measured by ELISA in HDL isolated from *Ldlr^−/−^* mice treated as described in Figure 1. Data are presented as mean ± SEM (N = 8 per group), *** p<0.001. One-way ANOVA with Bonferroni’s post-test. (B) Western blots of apoAI and MDA-apoAI in HDL isolated from plasma by immunoprecipitation using primary anti-apoAI antibody. *Ldlr^−/−^* mice were treated as described in Figure 1 and apoAI and MDA-apoAI from *Ldlr^−/−^* mice consuming a chow diet are included for comparison. (C) Quantitation using ImageJ software of the mean density ratio (arbitrary units) of MDA-apoAI to apoAI detected by Western blotting (B). (D) The HDL was isolated from the plasma of *Ldlr^−/−^* mice consuming a western diet for 16 weeks and treated with 2-HOBA or 4-HOBA or vehicle. Cholesterol enriched macrophages were incubated for 24h with HDL (25 μg protein/ml), and the % reduction in cellular cholesterol content measured. Data presented as mean ± SEM, N = 7 per group, * p<0.05, ** p<0.01, One-way ANOVA with Bonferroni’s post-test. (E) The MDA adducts were measured by ELISA in HDL isolated from control or FH subjects before and after LDL apheresis. N = 7 or 8, *** p<0.001, One-way ANOVA with Bonferroni’s post-test. (F) The MDA-Lysyl crosslink content in HDL from control or FH subjects (n=6 per group), p=0.02, Mann-Whitney test. (G) The capacity of HDL from control or FH subjects pre and post LDL apheresis (n=7 per group) to reduce the cholesterol content of *Apoe^−/−^* macrophages.

## Discussion

Oxidative stress-induced lipid peroxidation has been implicated in the development of atherosclerosis. Genetic defects and/or environmental factors cause an imbalance between oxidative stress and the ability of the body to counteract or detoxify the harmful effects of oxidation products ^1, 3, 34^. The large body of experimental evidence implicating an important role of lipid peroxidation in the pathogenesis of atherosclerosis previously had stimulated interest in the potential for antioxidants to prevent atherosclerotic cardiovascular disease. Although a few trials of dietary antioxidants in humans demonstrated reductions in atherosclerosis and cardiovascular events, the majority of large clinical outcomes trials with antioxidants have failed to show any benefit in terms of reduced cardiovascular events. Possible reasons for the failure of these trials to reduce cardiovascular events, include inadequate doses of antioxidants being used in the trials ^1, 16^ and the inhibition of normal ROS signaling that may be anti-atherogenic ^35^.

Peroxidation of lipids in tissues/cells or in blood produces a number of reactive lipid carbonyls and dicarbonyls including 4-hydroxynonenal, methylglyoxal, malondialdehyde, 4-oxo-nonenal, and isolevuglandins. These electrophiles can covalently bind to proteins, phospholipids, and DNA causing alterations in lipoprotein and cellular functions ^1, 10, 11^. Treatment with scavengers of reactive lipid carbonyl and dicarbonyl species represents a novel alternative therapeutic strategy that will decrease the adverse effects of a particular class of bioactive lipids without completely inhibiting the normal signaling mediated by ROS ^35^. A number of compounds with the potential to scavenge carbonyls have been identified, with individual compounds preferentially reacting with different classes of carbonyls so that the effectiveness of a scavenging compound in mitigating disease can serve as an indicator that their target class of carbonyl contributes to the disease process^35^. Previous studies found that scavengers of methylglyoxal and glyoxal, such as aminoguanidine and pyridoxamine, reduce atherosclerotic lesions in streptozotocin-treated *Apoe^−/−^* mice^36, 37^. Similarly, scavengers of α,β-unsaturated carbonyls (e.g. HNE and acrolein) such as carnosine and its derivatives, also reduce atherosclerosis in *Apoe^−/−^* mice or streptozotocin-treated *Apoe^−/−^* mice^38, 39, 40^. These previously tested scavenger compounds are poor in vivo scavengers of lipid dicarbonyls such as IsoLG and MDA^35^. Therefore, we sought to examine the potential of 2-HOBA, an effective scavenger of IsoLG and MDA, to prevent the development of atherosclerosis in *Ldlr^−/−^* mice.

We have recently reported that 2-HOBA can reduce isolevuglandin-mediated HDL modification and dysfunction ^41^. Our current studies are the first to examine the effects of dicarbonyl scavenging on atherosclerosis, and we demonstrate that the dicarbonyl scavenger, 2-HOBA, significantly reduces atherosclerosis development in the hypercholesterolemic *Ldlr^−/−^* mouse model (Figure 1). Importantly, our studies show that 2-HOBA treatment markedly improves features of the stability of the atherosclerotic plaque as evidenced by decreased necrosis and increased fibrous cap thickness and collagen content (Figure 3). Consistent with the proinflammatory effects of reactive dicarbonyls ^41^ and the impact on lesion necrosis, 2-HOBA reduced systemic inflammation by neutralizing reactive dicarbonyls (Figures 5 and 6). Furthermore, dicarbonyl scavenging reduced in vivo MDA modification of HDL, consistent with the notion that preventing dicarbonyl modification of HDL improves its net cholesterol efflux capacity (Figure 7). We previously showed that IsoLG modification increases in HDL from subjects with familial hypercholesterolemia ^41^, and this current study shows that MDA modification is similarly increased (Figure 7), suggesting these modifications contribute to the enhanced foam cell formation induced by FH-HDL (Figure 7). Taken together, dicarbonyl scavenging using 2-HOBA offers therapeutic potential in reducing atherosclerosis development and the risk of clinical events resulting from formation of vulnerable atherosclerotic plaques.

As our studies show that 2-HOBA reduces atherosclerosis development without decreasing plasma cholesterol levels (Figure 1), the atheroprotective effects of 2-HOBA are likely due to scavenging bioactive dicarbonyls. Consistent with this concept, the atherosclerotic lesion MDA- and IsoLG-lysyl adducts were decreased in 2-HOBA treated *Ldlr^−/−^* mice (Figure 2). That the effects of 2-HOBA are mediated by their action as dicarbonyl scavengers is further supported by the result that 4-HOBA, a geometric isomer of 2-HOBA, which is not an effective scavenger in vitro, is not atheroprotective and by the finding that MDA- and IsoLG-2-HOBA were abundantly formed versus −4-HOBA adducts (Supplemental Figures 5 and 6 and Table) in hypercholesterolemic *Ldlr^−/−^* mice. In addition, the levels of urine F_2_-isoprostanes were not significantly different between 2-HOBA and 4-HOBA treated *Ldlr^−/−^* mice suggesting that the atheroprotective effects are not via inhibition of lipid peroxidation or chelating metal ions (Supplemental Figure 7). A limitation to our study is the finding that 4-HOBA is cleared more rapidly compared to 2-HOBA in vivo, raising the possibility that our finding that 4-HOBA dicarbonyl adducts were very low to undetectable in vivo could in part be due to the lower concentrations of 4-HOBA in tissues. However, it is important to note that when macrophages were treated in vitro with ox-LDL in the presence of 2-HOBA or 4-HOBA, 2-HOBA-MDA adducts were readily detected, whereas 4-HOBA-MDA adducts were undetectable (Supplemental Figure 8), supporting the lack of reactivity of 4-HOBA with reactive dicarbonyls in biological systems. Nonetheless, our studies showing that atherosclerosis can be prevented by utilizing 2-HOBA to remove dicarbonyls strengthens the hypothesis that reactive dicarbonyls contribute to the pathogenesis of atherogenesis and raises the therapeutic potential of dicarbonyl scavenging. In this regard, we found that *Ldlr^−/−^* mice treated with 1g of 2-HOBA/L of water had plasma levels of 2-HOBA that were similar to humans receiving oral doses of 2-HOBA in our recent safety trial in humans ^24^.

HDL mediates a number of atheroprotective functions and evidence has mounted that markers of HDL dysfunction, such as impaired cholesterol efflux capacity, may be a better indicator of CAD risk than HDL-C levels ^1, 7, 42, 43, 44^. Patients with FH have previously been shown to have impaired HDL cholesterol efflux capacity, indicative of dysfunctional HDL ^45, 46^. Our studies show that consumption of a western diet by *Ldlr^−/−^* mice results in enhanced MDA-apoAI adduct formation (Figure 7), and that 2-HOBA treatment dramatically reduces modification of both apoAI and HDL with MDA. Similarly, FH patients had increased plasma levels of MDA-HDL adducts. In addition, in vitro modification of HDL with MDA resulted in decreased net cholesterol efflux capacity, similar to what we showed previously with IsoLG ^41^, and these effects which were observed with HDL containing MDA adducts in the same range as FH subjects and hypercholesterolemic mice (Figure 7 and Supplemental Figure 12). Our results do not agree with other studies showing that MDA modification of HDL does not significantly impact cholesterol efflux from cholesterol-enriched P388D_1_ macrophages, which may be due to differences in modification conditions or cell type ^32^. Our findings are consistent with studies by Shao and colleagues demonstrating that modification of lipid-free apoAI with MDA blocks ABCA1 mediated cholesterol efflux ^31^. In addition, studies have shown that long term cigarette smoking causes increased MDA-HDL adduct formation, and smoking cessation leads to improved HDL function with increased cholesterol efflux capacity ^47^. In line with these results, we found that HDL isolated from 2-HOBA versus vehicle and 4-HOBA treated mice has enhanced capacity to reduce cholesterol stores in macrophage foam cells (Figure 7). Furthermore, HDL from human subjects with FH had markedly increased MDA adducts and severely impaired ability to reduce macrophage cholesterol stores pre- and post-LDL apheresis (Figure 7). Thus, one of the atheroprotective mechanisms of 2-HOBA is likely through preventing formation of dicarbonyl adducts of HDL proteins, thereby preserving HDL net cholesterol efflux function. In addition to decreasing HDL oxidative modification, our studies show that 2-HOBA treatment decreases the in vivo MDA modification of of plasma LDL. Studies have shown that MDA modification of LDL promotes uptake via scavenger receptors resulting in foam cell formation and an inflammatory response ^48, 49^. The finding that incubation of macrophages with LDL from both 2-HOBA and 4-HOBA treated mice resulted in a similar cholesterol content is consistent with LDL, which is modified with sufficient amounts of MDA, being rapidly removed via scavenger receptors. However, studies have shown that neutralization of MDA-apoB adducts with antibodies greatly enhances atherosclerosis regression in human apoB100 transgenic *Ldlr^−/−^* mice ^50, 51^ making it likely that the decreased atherosclerosis with 2-HOBA treatment is also due in part to decreased dicarbonyl modification of apoB within the atherosclerotic lesion.

Evidence has mounted that increased oxidative stress in arterial intima cells is pivotal in inducing ER stress, inflammation, and cell death in atherogenesis ^52, 53^. In particular, efficient efferocytosis and limited cell death are critical to preventing the necrosis and excessive inflammation characteristic of the vulnerable plaque ^1, 52, 54^. Our results demonstrate that treatment with 2-HOBA promotes characteristics of more stable atherosclerotic plaques in *Ldlr^−/−^* mice (Figure 3). Consistent with this possibility, our study shows that 2-HOBA treatment decreased the atherosclerotic lesion MDA and IsoLG adduct content (Figure 2), supporting the ability of dicarbonyl scavenging in the arterial intima to limit oxidative stress induced inflammation, cell death, and destabilization of the plaque. Our studies show that scavenging of dicarbonyls with 2-HOBA in vitro limits oxidative stress induced apoptosis in both endothelial cells and macrophages (Figure 6). The decreased cell death is likely due in part to the greatly diminished inflammatory response to oxidative stress from dicarbonyl scavenging with 2-HOBA, as evidenced by the dramatic reductions in serum inflammatory cytokines including IL-1β (Figure 5). These results are particularly relevant given the recent results of the CANTOS trial showing that reducing inflammation with canakinumab, an IL-1β neutralizing monoclonal antibody, can reduce cardiovascular event rates in humans with prior MI and elevated hsCRP ^55^. Importantly, treatment with 2-HOBA did not impact levels of urinary prostaglandin metabolites of prostacyclin, thromboxane, PGE2 and PGD2, indicating that 2-HOBA does not result in significant inhibition of cyclooxygenase in mice in vivo (Supplemental Figure 10). Furthermore, 2-HOBA treatment maintained efficient efferocytosis and reduced the number of dead cells in the atherosclerotic lesions (Figure 4). As a result, dicarbonyl scavenging with 2-HOBA promoted features of stable plaques with decreased necrosis and enhanced collagen content and fibrous cap thickness (Figure 3). Hence, the ability of 2-HOBA to limit death and inflammation in arterial cells in response to oxidative stress and to promote efficient efferocytosis in the artery wall provides a novel atheroprotective mechanism whereby dicarbonyl scavenging promotes features of plaque stabilization and reduces atherosclerotic lesion formation. Our results are substantiated by recent studies demonstrating that *Ldlr^−/−^* mice expressing the single-chain variable fragment of E06 antibody to oxidized phospholipid have decreased atherosclerosis with stable plaque features including decreased necrosis and systemic inflammation ^56^, effects that are likely due in part to neutralization of esterified reactive dicarbonyls. Given the findings that significant residual inflammatory risk for CAD clinical events in humans independent of cholesterol lowering ^55, 57^, these studies highlight reactive dicarbonyls as a target to decrease this risk. The prevention of atherosclerotic lesion formation is clearly an important strategy for the prevention of cardiovascular events. A limitation of the current study is that we have not examined the impact of 2-HOBA as an intervention for established lesions. In this regard, our future studies will be directed at examining whether reactive dicarbonyl scavenging can remodel established atherosclerotic lesions in *Ldlr^−/−^* mice.

In conclusion, 2-HOBA treatment suppresses atherosclerosis development in hypercholesterolemic *Ldlr^−/−^* mice. The atheroprotective effects of 2-HOBA likely result from preventing dicarbonyl adduct formation with plasma apoproteins and intimal cellular components. Treatment with 2-HOBA decreased the formation of MDA-apoAI adducts thereby maintaining efficient HDL function. In addition, the prevention of MDA-apoB adducts decreases foam cell formation and inflammation. Finally, within the atherosclerotic lesion, dicarbonyl scavenging limited cell death, inflammation, and necrosis thereby effectively promoting characteristics of stable atherosclerotic plaques. As the atheroprotective effect of 2-HOBA treatment is independent of any action on serum cholesterol levels, 2-HOBA offers real therapeutic potential for decreasing the residual CAD risk that persists in patients treated with HMG-CoA reductase inhibitors.

## Materials and Methods

### Mice

*Ldlr^−/−^* and WT on C57BL/6 background mice were obtained from the Jackson Laboratory. Animal protocols were performed according to the regulations of Vanderbilt University’s Institutional Animal Care and Usage Committee. Mice were maintained on chow or a Western-type diet containing 21% milk fat and 0.15% cholesterol (Teklad). Eight week old, female *Ldlr^−/−^* mice on a chow diet were pretreated with vehicle alone (Water) or containing either 1 g/L of 4-HOBA or 1 g/L of 2-HOBA. 4-HOBA (as hydrochloride salt) was synthesized as previously described^21^. 2-HOBA (as the acetate salt, CAS 1206675-01-5) was manufactured by TSI Co., Ltd. (Missoula, MT) and obtained from Metabolic Technologies, Inc., Ames, IA^24^. A commercial production lot was used (Lot 16120312), and the purity of the commercial lot was verified to be > 99% via HPLC and NMR spectroscopy^24^. After two weeks, the mice continued to receive these treatments but were switched to a western diet for 16 weeks to induce hypercholesterolemia and atherosclerosis. Similarly, 12 week-old male *Ldlr^−/−^* mice were pretreated with vehicle alone (water) or containing 1 g/L of 2-HOBA for two weeks and were then switched to a western diet for 16 weeks to induce hypercholesterolemia and atherosclerosis, while continuing the treatment with 2-HOBA or water alone ^58, 59, 60^. Based on the average weight and daily consumption of water per mouse the estimated daily dosage with 1g/L of 2-HOBA is 200 mg/Kg. We did not observe differences in mouse mortality among the treatment groups. Eight week old, male *Ldlr^−/−^* mice were fed a western diet for 16 weeks and were continuously treated with water containing either 2-HOBA or 4-HOBA. Urine samples were collected using metabolic cages (2 mice in one cage) during 18 h after oral gavage with either 2-HOBA or 4-HOBA (5 mg each mouse).

### Cell Culture

Peritoneal macrophages were isolated from mice 72 hours post injection of 3% thioglycollate and maintained in DMEM plus 10% fetal bovine serum (FBS, Gibco) as previously described ^30^. Human aortic endothelial cells (HAECs) were obtained from Lonza and maintained in endothelial cell basal medium-2 plus 1% FBS and essential growth factors (Lonza).

### Plasma Lipids and Lipoprotein Distribution Analyses

The mice were fasted for 6 hours, and plasma total cholesterol and triglycerides were measured by enzymatic methods using the reagents from Cliniqa (San-Macros, CA). Fast performance liquid chromatography (FPLC) was performed on an HPLC system model 600 (Waters, Milford, MA) using a Superose 6 column (Pharmacia, Piscataway, NJ).

### HDL Isolation from Mouse Plasma and Measurement of HDL Capacity to Reduce Macrophage Cholesterol

HDL was isolated from mouse plasma using HDL Purification Kit (Cell BioLabs, Inc.) following the manufacturer’s protocol. Briefly, apoB containing lipoproteins and HDL were sequentially precipitated with dextran sulfate. The HDL was then resuspended and washed. After removing the dextran sulfate, the HDL was dialyzed against PBS. To measure the capacity of the HDL to reduce macrophage cholesterol, *Apoe^−/−^* macrophages were cholesterol enriched by incubation for 48h in DMEM containing 100 μg protein/ml of acetylated LDL. The cells were then washed, and incubated for 24h in DMEM alone or with 25 μg HDL protein/ml. Cellular cholesterol was measured before and after incubation with HDL using an enzymatic cholesterol assay as described ^61^.

### Human Blood Collection and Measurement of MDA-LDL, MDA-HDL, and MDA-ApoAI

The study was approved by the Vanderbilt University Institutional Review Board (IRB), and all participants gave their written informed consent. The human blood samples from patients with severe FH, who were undergoing LDL apheresis, and healthy controls were obtained using an IRB approved protocol. HDL and LDL were prepared from serum by Lipoprotein Purification Kits (Cell BioLabs, Inc.). Sandwich ELISA was used to measure plasma MDA-LDL and MDA-HDL levels following the manufacturer’s instructions (Cell BioLabs, Inc.). Briefly, isolated LDL or HDL samples and MDA-Lipoprotein standards were added onto anti-MDA coated plates, and, after blocking, the samples were incubated with biotinylated anti-apoB or anti-ApoAI primary antibody. The samples were then incubated for 1h with streptavidin-enzyme conjugate and 15 min with substrate solution. After stopping the reaction, the O.D. was measured at 450 nm wavelength. MDA-ApoAI was detected in mouse plasma by immunoprecipitation of ApoAI and western blotting. Briefly, 50 μl of mouse plasma was prepared with 450 μL of IP Lysis Buffer (Pierce) plus 0.5% protease inhibitor mixture (Sigma), and immunoprecipitated with 10 μg of polyclonal antibody against mouse ApoAI (Novus). Then 25 μL of magnetic beads (Invitrogen) was added, and the mixture was incubated for 1h at 4°C with rotation. The magnetic beads were then collected, washed three times, and SDS-PAGE sample buffer with β-mercaptoethanol was added to the beads. After incubation at 70°C for 5 min, a magnetic field was applied to the Magnetic Separation Rack (New England), and the supernatant was used for detecting mouse ApoAI or MDA. For Western blotting, 30-60 μg of proteins was resolved by NuPAGE Bis-Tris electrophoresis (Invitrogen), and transferred onto nitrocellulose membranes (Amersham Bioscience). Membranes were probed with primary rabbit antibodies specific for ApoAI (Novus NB600-609) or MDA-BSA (Abcam cat# ab6463) and fluorescent tagged IRDye 680 (LI-COR) secondary antibody. Proteins were visualized and quantitated by Odyssey 3.0 Quantification software (LI-COR).

### Modification of HDL and LDL with MDA

MDA was prepared immediately before use by rapid acid hydrolysis of maloncarbonyl bis- (dimethylacetal) as described ^31^. Briefly, 20 μL of 1 M HCl was added to 200 μL of maloncarbonyl bis-(dimethylacetal), and the mixture was incubated for 45 min at room temperature. The MDA concentration was determined by absorbance at 245 nm, using the coefficient factor 13, 700 M^−1^ cm^−1^. HDL (10mg of protein /mL) and increasing doses of MDA (0, 0.125 mM, 0.25 mM, 0.5 mM, 1 mM) were incubated at 37 °C for 24 h in 50 mM sodium phosphate buffer (pH7.4) containing DTPA 100 μM. Reactions were initiated by adding MDA and stopped by dialysis of samples against PBS at 4 °C. LDL (5 mg/mL) was modified in vitro with MDA (10 mM) in the presence of vehicle alone or with 2-HOBA at 37°C for 24 h in 50 mM sodium phosphate buffer (pH7.4) containing DTPA 100 μM. Reactions were initiated by adding MDA and stopped by dialysis of samples against PBS at 4 °C. The LDL samples were incubated for 24h with macrophages and the cholesterol content of the cells was measured using an enzymatic cholesterol assay as described ^61^.

### Atherosclerosis Analyses and Cross-section Immunofluorescence Staining

The extent of atherosclerosis was examined both Oil-Red-O-stained cross-sections of the proximal aorta and by *en face* analysis ^30^. Briefly, cryosections of 10-micron thickness were cut from the region of the proximal aorta starting from the end of the aortic sinus and for 300 μm distally, according to the method of Paigen et al. ^62^. The Oil red-O staining of 15 serial sections from the root to ascending aortic region were used to quantify the Oil red-O-positive staining area per mouse. The mean from the 15 serial sections was applied for the aortic root atherosclerotic lesion size per mouse using the KS300 imaging system (Kontron Elektronik GmbH) as described ^63, 64, 65^. All other stains were done using sections that were 40 to 60 μm distal of the aortic sinus. For each mouse, 4 sections were stained and quantitation was done on the entire cross section of all 4 sections. For immunofluorescence staining, 5 μm cross-sections of the proximal aorta were fixed in cold acetone (Sigma), blocked in Background Buster (Innovex), incubated with indicated primary antibodies (MDA and CD68) at 4°C for overnight. After incubation with fluorescent labeled secondary antibodies at 37C for 1 hour, the nucleus was counterstained with Hoechst. Images were captured with a fluorescence microscope (Olympus IX81) and SlideBook 6 (Intelligent-Image) software and quantitated using ImageJ software (NIH) ^66^.

### In vitro Cellular Apoptosis and Analysis of Lesion Apoptosis and Efferocytosis

Cell apoptosis was induced as indicated and detected by fluorescent labeled Annexin V staining and quantitated by either Flow Cytometry (BD 5 LSRII) or counting Annexin V positive cells in images captured under a fluorescent microscope. The apoptotic cells in atherosclerotic lesions were measured by TUNEL staining of cross-sections of atherosclerotic proximal aortas as previously described ^30^. The TUNEL positive cells not associated with live macrophages were considered free apoptotic cells and macrophage-associated apoptotic cells were considered phagocytosed as a measure of lesion efferocytosis as previously described ^30^.

### Masson’s Trichrome Staining

Masson’s Trichrome Staining was applied for measurement of atherosclerotic lesion collagen content, fibrous cap thickness and necrotic core size following the manufacture’s instructions (Sigma) and as previously described ^30^. Briefly, 5 μm cross-sections of proximal atherosclerotic aorta root were fixed with Bouin’s solution, stained with hematoxylin for nuclei (black) and biebrich scarlet and phosphotungstic/phosphomolybdic acid for cytoplasm (red), and aniline blue for collagen (blue). Images were captured and analyzed for collagen content, atherosclerotic cap thickness and necrotic core by ImageJ software as described previously ^30^. The necrotic area is normalized to the total lesion area and is expressed as the % necrotic area.

### RNA Isolation and Real-Time RT-PCR

Total RNA was extracted and purified using Aurum Total RNA kit (Bio-Rad) according to the manufacturer’s protocol. Complementary DNA was synthesized with iScript reverse transcriptase (Bio-Rad). Relative quantitation of the target mRNA was performed using specific primers, SYBR probe (Bio-Rad), and iTaqDNA polymerase (Bio-Rad) on IQ5 Thermocylcer (Bio-Rad) and normalized with 18S, as described earlier. 18S, IL-1β and TNF-α primers used were as described earlier ^67^.

### Liquid chromatography-mass spectrometry analysis of urinary prostaglandin metabolites

Concentrations of PGE-M, tetranor PGD-M, 11-dehydro-TxB^2^ (TxB-M) and PGI-M in urine were measured in the Eicosanoid Core Laboratory at Vanderbilt University Medical Center. Urine (1mL) was acidified to pH 3 with HCl. [^2^H_4_]-2,3-dinor-6-keto-PGF1a (internal standard for PGI-M quantification) and [^2^H_4_]-11-dehydro-TxB_2_ were added, and the sample was treated with methyloxime HCl to convert analytes to the O-methyloxime derivative. The derivatized analytes were extracted using a C-18 Sep-Pak (Waters Corp. Milford, MA USA) and eluted with ethyl acetate as previously described ^68^. A [^2^H_6_]-O-methyloxime PGE-M deuterated internal standard was then added for PGE-M and PGD-M quantification. The sample was dried under a stream of dry nitrogen at 37°C and then reconstituted in 75 μL mobile phase A for LC/MS analysis.

LC was performed on a 2.0 × 50 mm, 1.7 μm particle Acquity BEH C18 column (Waters Corporation, Milford, MA, USA) using a Waters Acquity UPLC. Mobile phase A was 95:4.9:0.1 (v/v/v) 5 mM ammonium acetate:acetonitrile:acetic acid, and mobile phase B was 10.0:89.9:0.1 (v/v/v) 5 mM ammonium acetate:acetonitrile:acetic acid. Samples were separated by a gradient of 85–5% of mobile phase A over 14 min at a flow rate of 375 μl/min prior to delivery to a SCIEX 6500+ QTrap mass spectrometer.

Urinary creatinine levels are measured using a test kit from Enzo Life Sciences. The urinary metabolite levels in each sample are normalized using the urinary creatinine level of the sample and expressed in ng/mg creatinine.

### Measurement of IsoLG-Lys in aorta

Isolation and LC/MS measurement of isolevuglandin-lysyl-lactam (IsoLG-Lys) adducts from aorta of 2-HOBA and 4-HOBA treated *Ldlr^−/−^* mice were performed using a Waters Xevo-TQ-Smicro triple quadrupole mass spectrometer as previously described ^69^.

### Detection of IsoLG adducts of 2-HOBA

To generate an internal standard for quantitation, 10 molar equivalents of the heavy isotope labeled 2-HOBA, [^2^H_4_]2-HOBA, was reacted with synthetic IsoLG^69^ overnight in 1 mM triethylammonium acetate buffer to form IsoLG-2-HOBA adducts, and the adducts separated from unreacted [^2^H_4_]2-HOBA and IsoLG by solid phase extraction (Oasis HLB). The isolated reaction products of IsoLG-2-HOBA were scanned by mass spectrometer (Waters Xevo-TQ-Smicro triple quadrupole MS) operating in limited mass scanning mode to identify major products. Additionally, precursor scanning with the product ion set at m/z 111.1 was used to confirm that the detected products were [^2^H_4_]2-HOBA adducts. Both methods showed that the primary adduct present in the purified IsoLG-[^2^H_4_]2-HOBA internal standard mixture was the IsoLG-[2H4]2-HOBA hydroxylactam adduct, although other adducts including pyrrole, lactam, and the anhydro-species of each of these adducts were also present. Similar species were seen when IsoLG was reacted with non-labeled 2-HOBA and precursor scanning using product ion m/z 107.1 To identify potential 2-HOBA adducts in tissue of treated animals, we first generated a list of 18 probable IsoLG-HOBA species [pyrrole, lactam, hydroxylactam based on the in vitro reactions of IsoLG and 2-HOBA and then the anhydro-, dinor-, dinor/anhydro-, tetranor-, and keto- (from oxidation of hydroxyl group) metabolites of each of these three adducts based on previous metabolism studies with prostaglandins and isoprostanes]. We then analyzed liver homogenate from a 2-HOBA treated mouse using LC/MS with the mass spectrometer operating in positive ion precursor scanning mode and the product ion set to m/z 107.1 and collision energy at 20eV and looked for the presence of any of these precursor ions. Based on these data, we identified three potential metabolites: M1 precursor ion m/z 438.3, which mass is consistent with either the keto-pyrrole adduct or the anhydro-lactam adduct (both have identical mass). M2 m/z 440.3, which mass is consistent with the pyrrole adduct, and M3 m/z 454.3 which mass is consistent with the anhydro-hydroxylactam adduct or the keto-lactam adduct. When then sought to quantify the amount of the putative IsoLG-HOBA adducts in heart and liver samples as there was not sufficient aorta sample remaining from other analysis available to do this analysis.

For these experiments, liver or heart samples from *Ldlr^−/−^* mice treated with 2-HOBA or 4-HOBA were homogenized in 0.5 M Tris buffer solution pH 7.5 containing mixture of antioxidants (pyridoxamine, indomethacin, BHT, TCEP). Total amount of protein in homogenate was determined for normalization. 1 pmol IsoLG-[^2^H_4_]2-HOBA was then added to each homogenate sample as internal standard, the HOBA adducts extracted with ethyl acetate, dried, dissolved in solvent 1 (water with 0.1% acetic acid) and analyzed by LC/MS using Waters Xevo-TQ-Smicro triple quadrupole mass spectrometer operating in positive ion multiple reaction monitoring (MRM) mode, monitoring the following transitions: m/z 438.3→107.1@20eV for M1; m/z 440→107.1@20eV for M2; m/z 454→107.1@20eV for M3; and m/z 476.3→111.1@20eV for IsoLG-[^2^H_4_]2-HOBA hydroxylactam. Desolvation temperature: 500°C; source temperature: 150°C; capillary voltage: 5 kV, cone voltage: 5 V; cone gas flow 1 L/h; desolvation gas flow 1000 L/h. HPLC condition were as follows: Solvent 1: water with 0.1% acetic acid; Solvent 2, methanol with 0.1% acetic acid; column: Phenomenex Kinetex C8 50 × 2.1 mm 2.6u 100A, flow rate: 0.4 mL/min; gradient: starting condition 10% B with gradient ramp to 100% B over 3.5 min, hold for 0.5 min, and return to starting conditions over 0.5 min. Abundance for each metabolite was calculated based on the ratio of peak height versus that of internal standard.

### Analysis of dilysyl-MDA crosslinks by LC/ESI/MS/MS

Samples (around 1 mg of protein) were digested with proteases as previously described for lysyl-lactam adducts ^70^. Five nanograms of ^13^C_6_-dilysyl-MDA crosslink standard were added to each cell sample and dilysyl-MDA crosslinks were purified as previously described ^71^. The dilysyl-MDA crosslink was quantified by isotopic dilution by LC-ESI/MS/MS as previously described ^71^.

### LC/MS/MS quantification of scavenger-MDA adducts

The scavenger-MDA adducts were extracted, (1) from homogenate of tissue (equivalent of 30 mg) or (2) from cells (1 ml), three times with 500 μl of ethyl acetate. The extract was dried down, resuspended in 100 μl of ACN-water (1:1, v/v with 0.1% formic acid), vortexed, and filtered through a 0.22 μm spin X column. The reactions were analyzed by LC-ESI/MS/MS using the column a Phenomenex Kinetex column at a flow rate of 0.1 ml/min. The gradient consisted of Solvent A, water with 0.2% formic acid and solvent B, acetonitrile with 0.2% formic acid. The gradient was as follows: 0-2 min 99.9% A, 2-9 min 99.9 - 0.1% A, 9 - 12min 99.9% B. The mass spectrometer was operated in the positive ion mode, and the spray voltage was maintained at 5,000 V. Nitrogen was used for the sheath gas and auxiliary gas at pressures of 30 and 5 arbitrary units, respectively. The optimized skimmer offset was set at 10, capillary temperature was 300°C, and the tube lens voltage was specific for each compound. SRM of specific transition ions for the precursor ions at m/z 178 → 107 (propenal-HOBA adduct).

### Statistics

Data are presented as mean ± SEM. The normality of the sample populations was examined by the Kolmogorov-Smirnov test, then differences between mean values were determined by oneway ANOVA (Bonferroni’s post-test), Kruskal-Wallis test (Bunn’s multiple comparison), Mann-Whitney test, and Student’s t-test using GraphPad PRISM. Significance was set for *p* < 0.05.

## Data Availability

The data generated from current studies are available from the corresponding author on reasonable request. Further information is also available in the supplemental figure and table that are linked to this paper within the Nature Reporting Summary system.

## Acknowledgments and Sources of Funding

The analyses of urinary prostaglandin metabolites were performed in the Vanderbilt University Eicosanoid Core Laboratory. 2-HOBA was obtained from Metabolic Technologies, Inc., Ames, IA. This work was supported by National Institutes of Health Grants: HL116263 and DK59637 (Lipid, Lipoprotein and Atherosclerosis Core of the Vanderbilt Mouse Metabolic Phenotype Centers).

## Authors’ contribution

H.T. designed and performed key experiments, acquired and analyzed data, wrote manuscript; J.H. designed and performed parts of experiments, acquired and analyzed data; P.G.Y. designed research, analyzed data, wrote manuscript; J. L. B., Y.Z., L.D., V. Y., and I.Z.I. conducted parts of the experiments; O.B., V.A., J.A.O., L.J.R.II, S.S.D.. designed and analyzed data and modified manuscript; M. F. L. designed research, analyzed data, wrote manuscript.

Huan Tao and Jiansheng Huang made equal contribution

