## Supplemental Figures for "Scavenging of reactive dicarbonyls with 2-hydroxybenzylamine reduces atherosclerosis in hypercholesterolemic *Ldlr^−/−^* mice"

Supplemental Figure 1

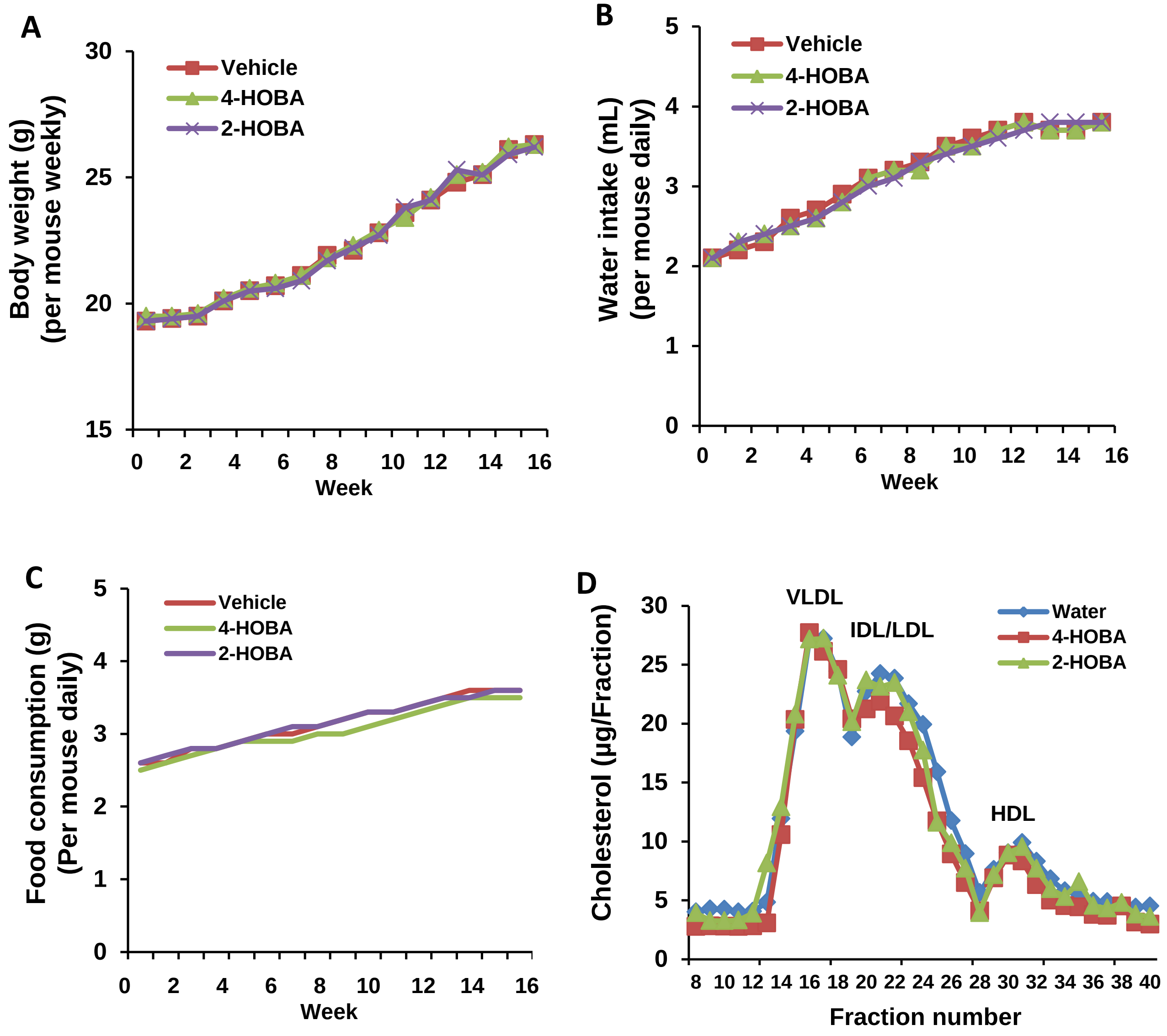


Supplemental Figure 1. 2-HOBA does not impact body weight, water intake, food consumption or lipoprotein profiles in hypercholesterolemic *Ldlr*$^{-/-}$ mice. The body weight (A), water intake (B), and diet consumption (C) were measured in the *Ldlr*$^{-/-}$ mice consuming a Western diet for 16 weeks and treated with 1 g/L 2-HOBA, 4-HOBA, or vehicle. (D) The plasma was pooled from hypercholesterolemic *Ldlr*$^{-/-}$ mice (4 mice/group) that were fasted for 6 hours. Fast performance liquid chromatography (FPLC) was performed using a Superose 6 column. Total cholesterol was measured by an enzymatic assay and the average is shown for two pooled plasma samples per group of mice.

Supplemental Figure 2

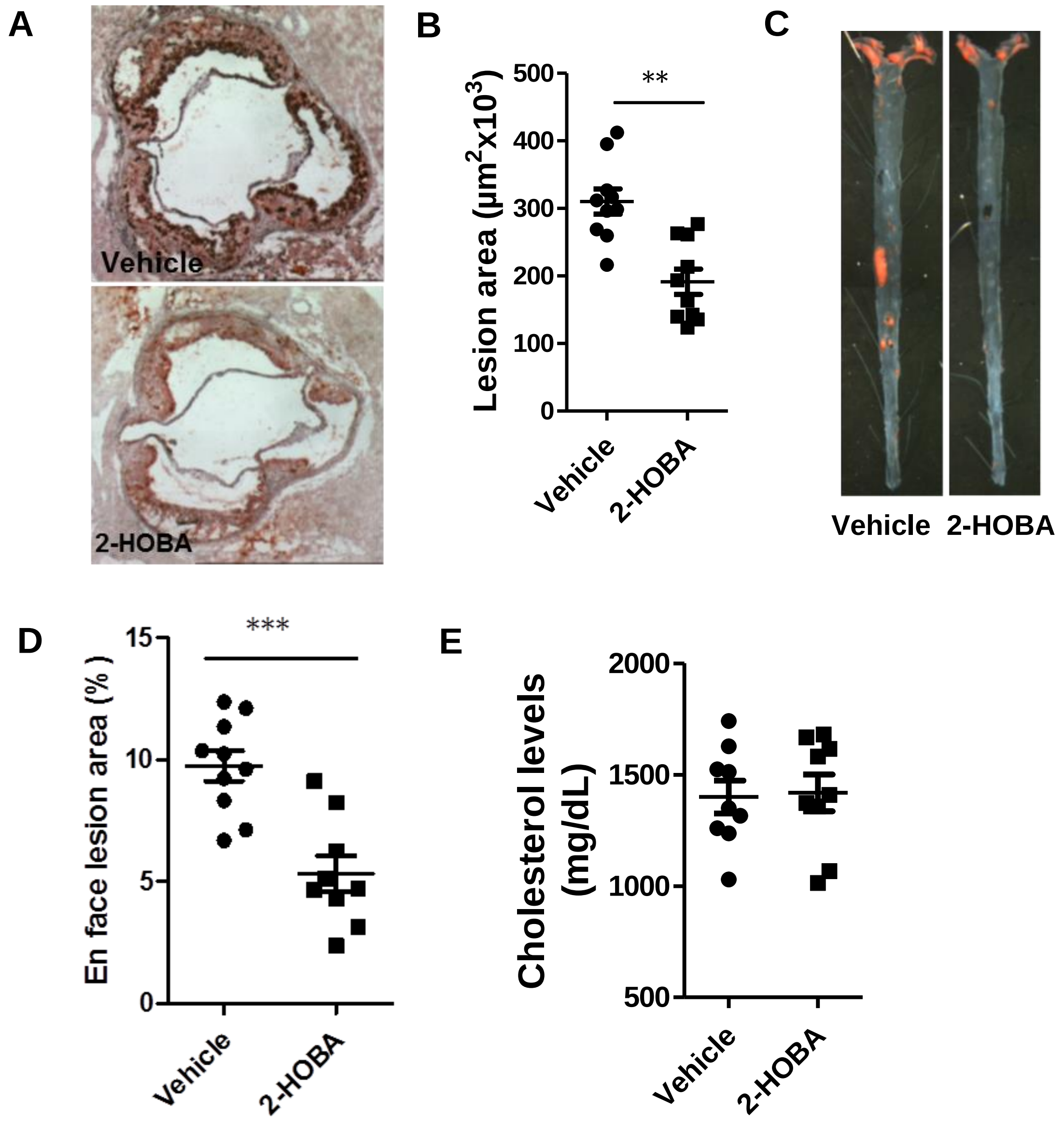


Supplemental Figure 2. 2-HOBA reduces atherosclerotic lesions in the hypercholesterolemic male *Ldlr*$^{-/-}$ mice. 12-week old male *Ldlr*$^{-/-}$ mice were pretreated with 1 g/L 2-HOBA or vehicle (water) for 2 weeks and then the treatment was continued for 16 weeks during which the mice were fed a Western diet. (A) Representative images show Red-Oil-O staing in the proximal aorta root sections. (B) Quantitation of the mean Oil-Red-O stainable lesion area in aortic root sections. (C) Representative images show Red-Oil-O staining in open-pinned aortas and (D) the quantitation of *en face* lesion area. (E) 2-HOBA does not affect the cholesterol levels of male *Ldlr*$^{-/-}$ mice. N =9 or 10 per group, ** $p<0.05$, . *** $p<0.001$, Student *t* test.

Supplemental Figure 3

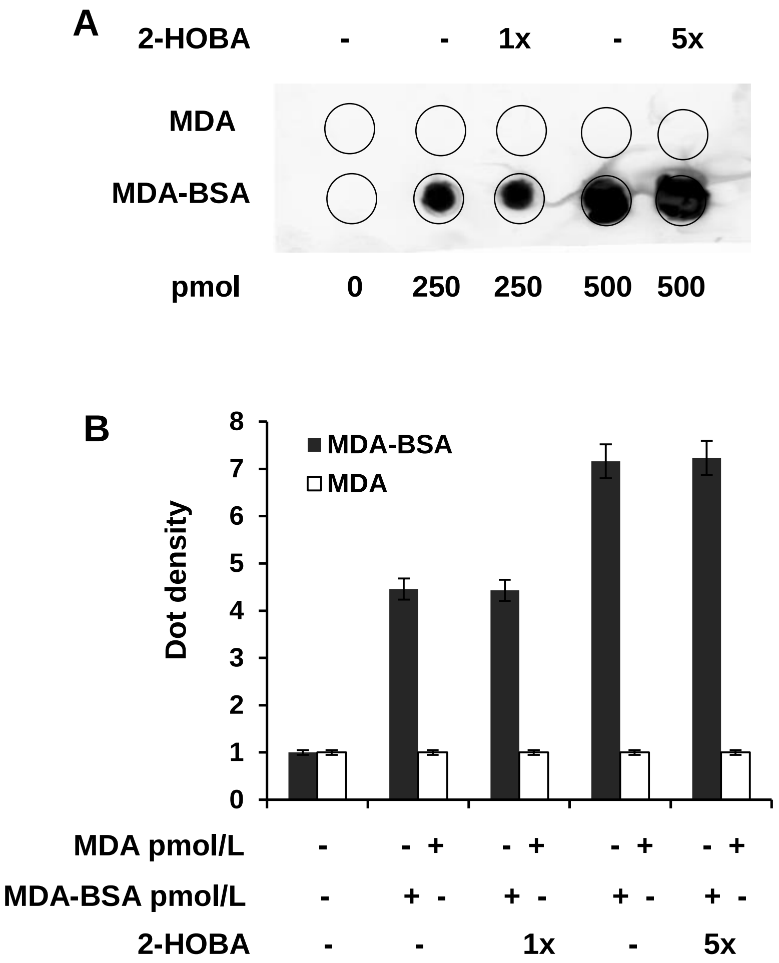


Supplemental Figure 3. 2-HOBA does not impact anti-MDA antibody interaction with MDA-BSA. A series of doses of MDA-BSA or MDA alone were incubated with 1x or 5x 2-HOBA. Then 2 µl of each sample was loaded onto HyBond-C membrane, and incubated with the blocking buffer, primary anti-MDA antibody and fluorescent seconary antibody after vigorous washing. The image was captured by the Odyssey system (A) and quantitated by ImageJ software (B).

Supplemental Figure 4

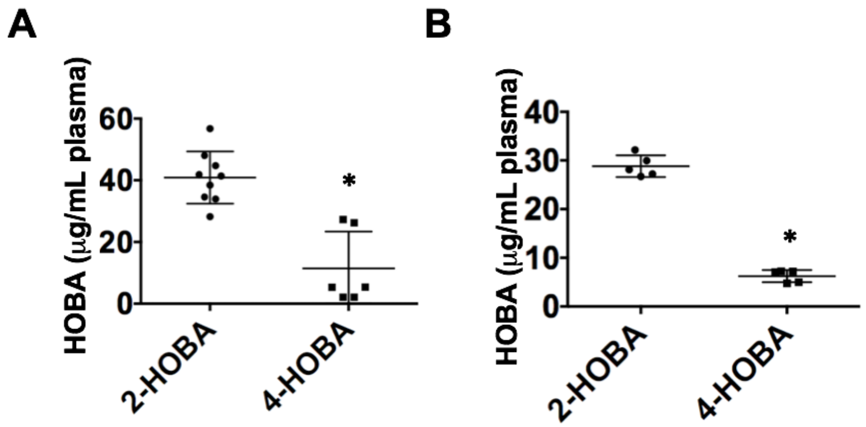


Supplemental Figure 4. The concentration of 2-HOBA or 4-HOBA was measured in plasma from mice. A. Eight week old male *Ldlr*$^{-/-}$ mice were fed WD for 16 weeks and were continuously treated with water containing either 2-HOBA (n=9) or 4-HOBA (n=6). Plasma was collected 30min after oral gavage of mice with either 2-HOBA or 4-HOBA (5mg each mouse). B. Plasma samples were collected from male C57BL6 mice on a chow diet 30min after intraperitoneal injection of 2-HOBA (n=5) or 4-HOBA (n=5). The concentration of 2-HOBA or 4-HOBA was measured in the plasma using LC-MS/MS as described in the supplemental methods. (Mann-Whitney test, * indicates $p<0.05$)

Supplemental Figure 5 and Table

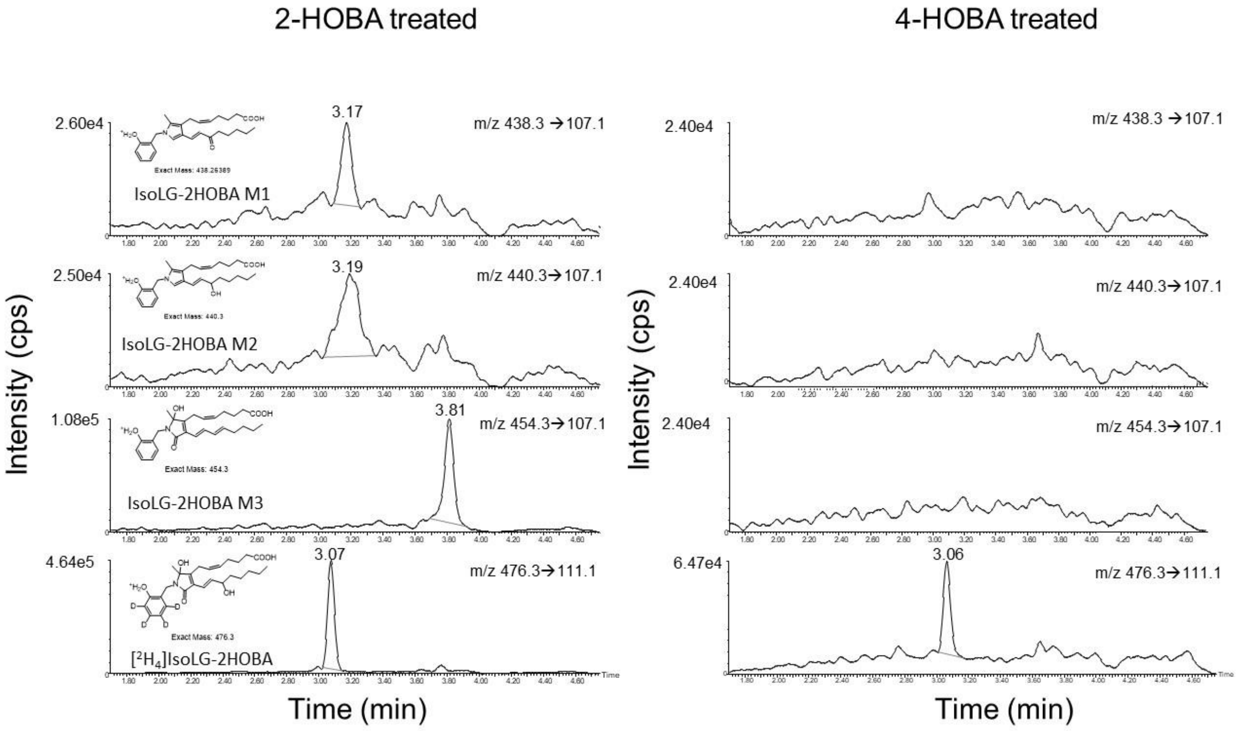


Supplemental Figure 5. Detection of metabolites of isolevuglandin modified 2-HOBA (IsoLG-2-HOBA) in liver of 2-HOBA treated *Ldlr*$^{-/-}$ mice. Putative metabolites were identified as described in supplemental methods. Representative chromatographs for livers from mice treated with 2-HOBA (left) and 4-HOBA (right) are shown for the three most abundant IsoLG-HOBA metabolites (three upper panels) and the internal standard (lower panel). One potential structure of each metabolite is shown on the left of the chromatograph.

Supplemental Table

| **Supplemental Table** | | |
|---|---|---|
| analytes | 2-HOBA treated | 4-HOBA treated |
| Liver (nmol/kg) | | |
| IsoLG-HOBA-M1 | 0.88±0.14 | ND |
| IsoLG-HOBA-M2 | 0.82±0.12 | ND |
| IsoLG-HOBA-M3 | 4.17±1.85 | ND |
| Heart (nmol/kg) | | |
| IsoLG-HOBA-M1 | 0.74±0.21 | ND |
| IsoLG-HOBA-M2 | 0.57±0.35 | ND |
| IsoLG-HOBA-M3 | 0.24±0.17 | ND |
| ND = not detected | | |

Supplemental Table: Levels of IsoLG-HOBA metabolites in liver and hearts of *Ldlr*$^{-/-}$ fed a Western diet for 16 weeks and continuously treated with water containing either 2-HOBA or 4-HOBA. Structures for metabolites 1-3 (M1, M2, M3) are shown in Supplemental Figure 6. No signal for IsoLG-HOBA metabolites were detected in mice treated with 4-HOBA. Livers and hearts from five mice for each group were analyzed and the mean and standard error of the mean is shown.

Supplemental Figure 6

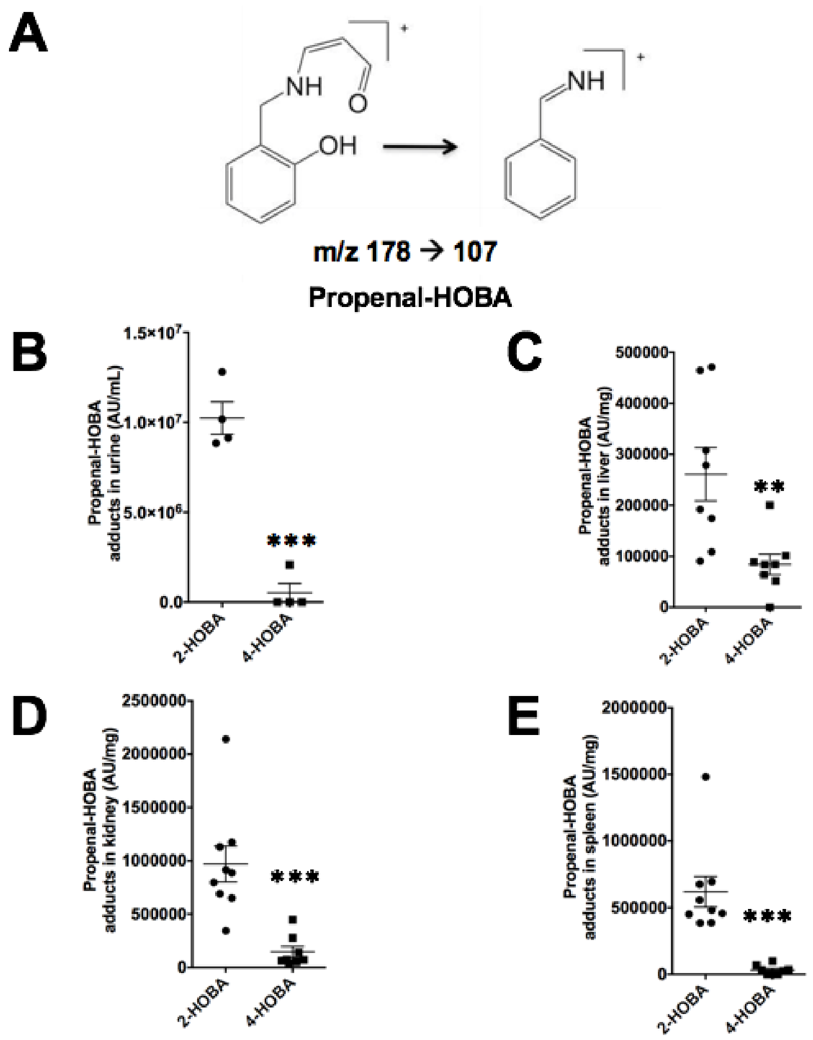


Supplemental Figure 6. MDA-2-HOBA adducts were more readily formed in vivo than MDA-4-HOBA adducts. Urine samples were collected after oral gavage of male *Ldlr*$^{-/-}$ mice on a WD with either 2-HOBA or 4-HOBA. HOBA-propenal adducts in urine (B), HOBA-propenal adducts in liver (C), HOBA-propenal adducts in kidney (D) and HOBA-propenal adducts in spleen (E) were measured in *Ldlr*$^{-/-}$ mice using LC-MS/MS as described in supplemental methods (Mann-Whitney test, ** indicates $p<0.01$ and *** indicates $p<0.001$).

Supplemental Figure 7

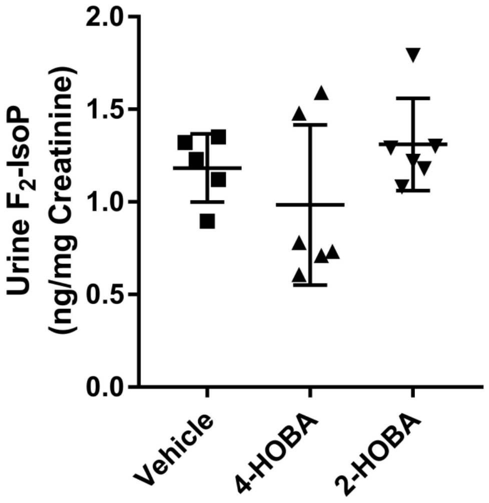


Supplemental Figure 7. 2-HOBA does not impact urine F2-IsoP in hypercholesterolemic *Ldlr*$^{-/-}$ mice. The urine F2-IsoP levels were measured by LC/MS/MS from *Ldlr*$^{-/-}$ mice consuming a western diet for 16 weeks and treated with 1 g/L 2-HOBA, 4-HOBA, or vehicle. N = 5 or 6 per group, p = 0.43, Kruskal-Wallis test. Urinary creatinine levels were measured for normalization.

Supplemental Figure 8

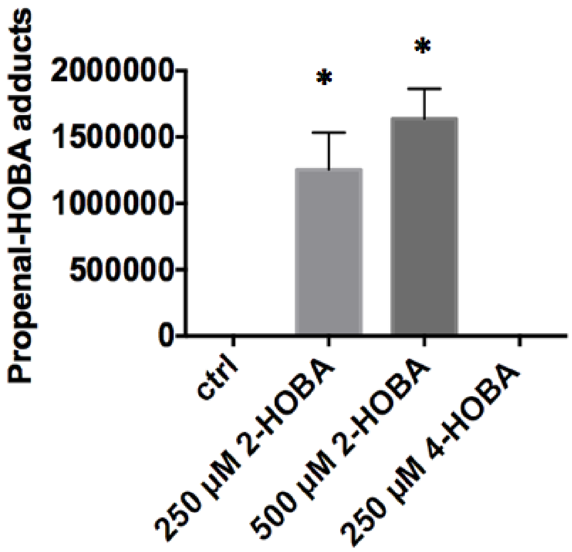


Supplemental Figure 8. Treatment of peritoneal macropahges with 2-HOBA increases the propenal-HOBA adducts levels. Peritoneal macrophages were isolated from C57BL6 mice and incubated with 50ug/mL ox-LDL, and treated with either 2-HOBA or 4-HOBA for 24h. Cell samples were collected and the HOBA-MDA adducts were measured using LC-MS/MS as described in supplemental methods. (One-way ANOVA with Bonferroni's Post-test).

Supplemental Figure 9

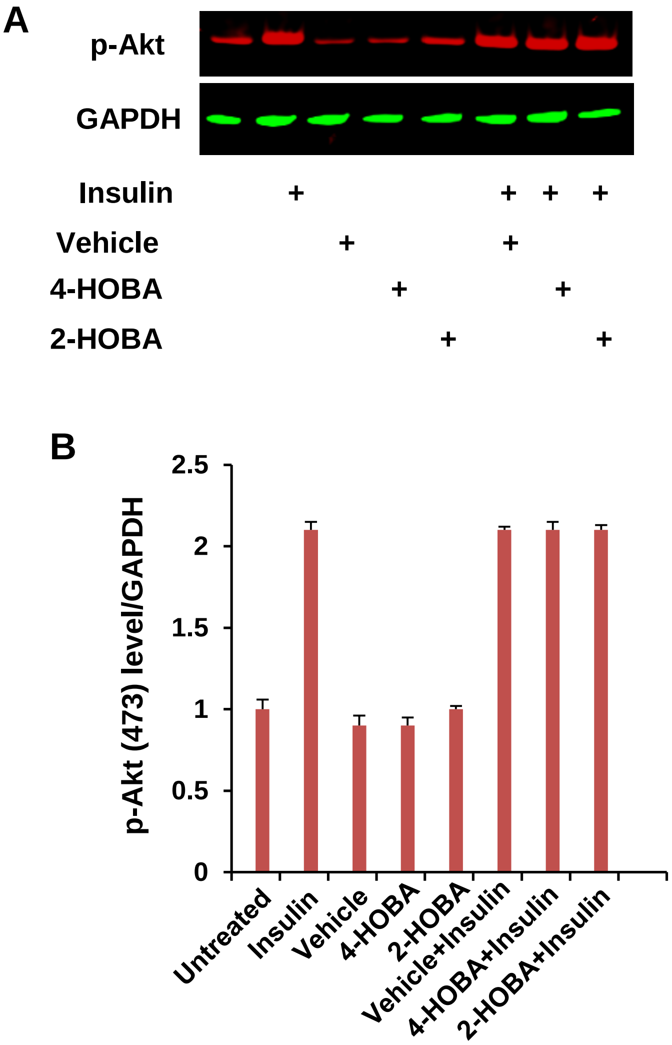


Supplemental Figure 9. 2-HOBA does not influence Akt signaling in macrophages. WT macrophages were treated with or without vehicle (water), 250 µM 4-HOBA or 2-HOBA for 1 hour, and then incubated with or without 100 nM insulin as indicated for 15 min. Phospho-Akt (S473) and GAPDH were detected by Western Blotting (A). The band density was quantitated by ImageJ software (B). Two independent experiments were performed.

Supplemental Figure 10

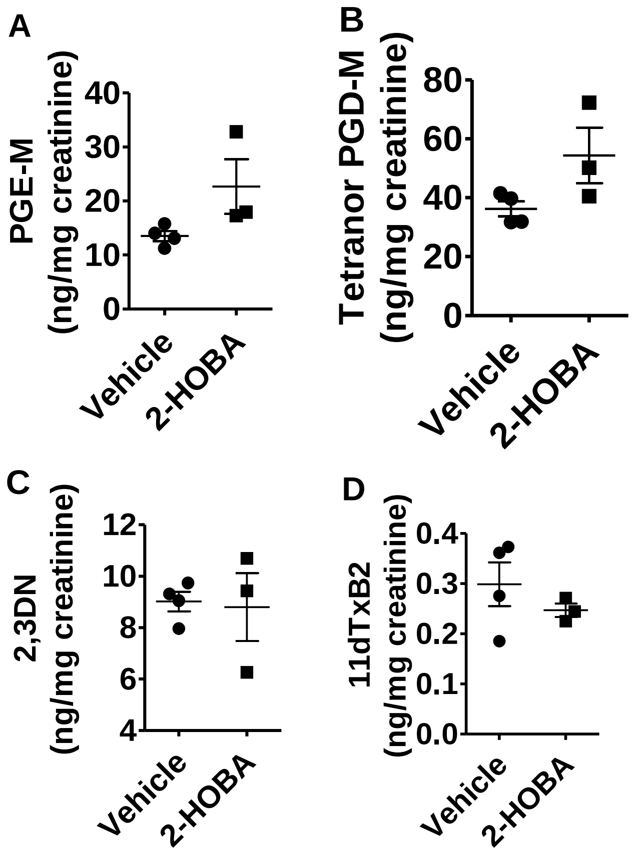


Supplemental Figure 10. Effect of 2-HOBA on prostaglandin metabolites. The urine samples were collected in metabolic cages with 2 mice per cage after 12 weeks of treatment with 2-HOBA or water. The contents of PGE-M (A), tetranor PGD-M (B), 2,3-dinor-6-keto-PGF1 (C) and 11-dehydro TxB2 (D) were analyzed by LC/MS/MS.

Supplemental Figure 11.

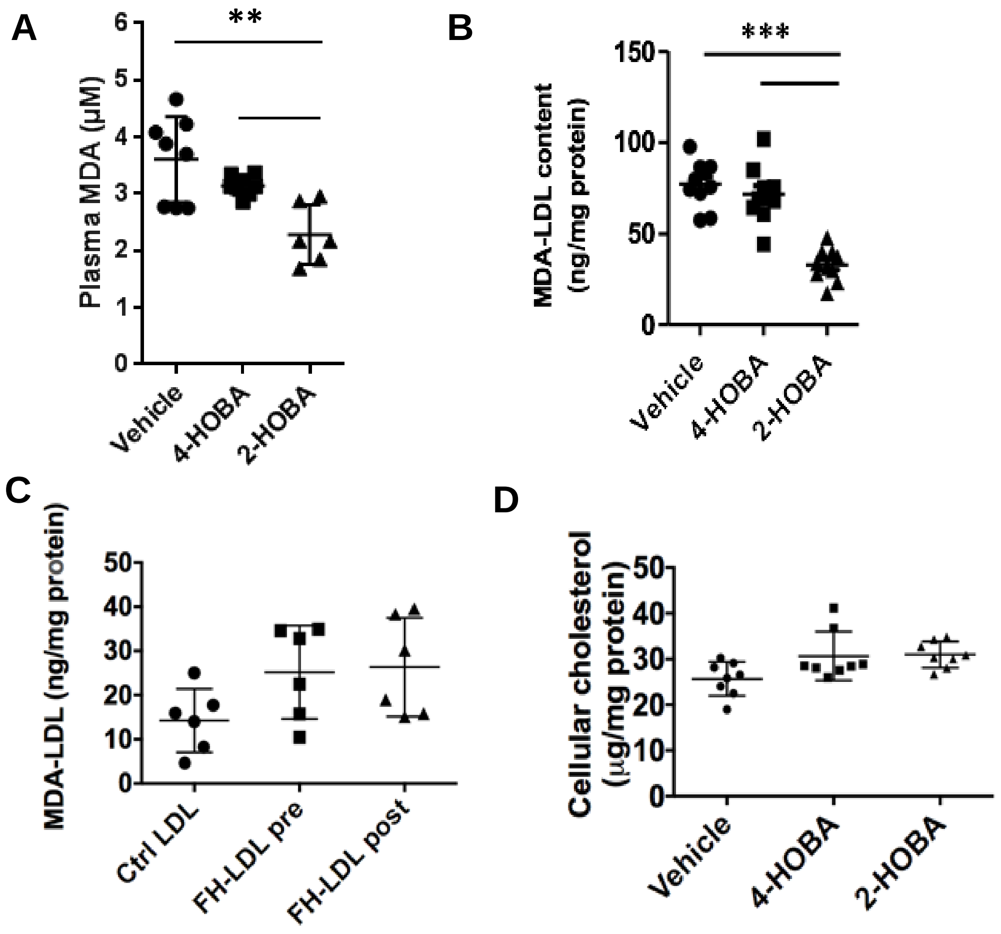


Supplemental Figure 11. Effects of 2-HOBA on plasma and LDL MDA adducts in hypercholesterolemic *Ldlr*$^{-/-}$ mice. (A) The MDA content in plasma from *Ldlr*$^{-/-}$ mice consuming a western diet for 16 weeks and treated with 2-HOBA, 4-HOBA, or vehicle was measured by TBARS Assay(**p*<0.05, ***p*<0.01). (B) The levels of MDA adducts were measured from LDL isolated from the *Ldlr*$^{-/-}$ mice by ELISA. N = 10 per group, *** p<0.001. (C) LDL was isolated from control and FH subjects (n=6) and the MDA adduct content was measured by ELISA. (D) LDL was isolated from 2-HOBA, 4-HOBA, or vehicle treated hypercholesterolemic *Ldlr*$^{-/-}$ mice. WT peritoneal macrophages were incubated for 24 hrs with the LDL and the cellular cholesterol content was measured as described in methods. (A-D) One-way ANOVA with Bonferroni's Post-test).

Supplemental Figure 12

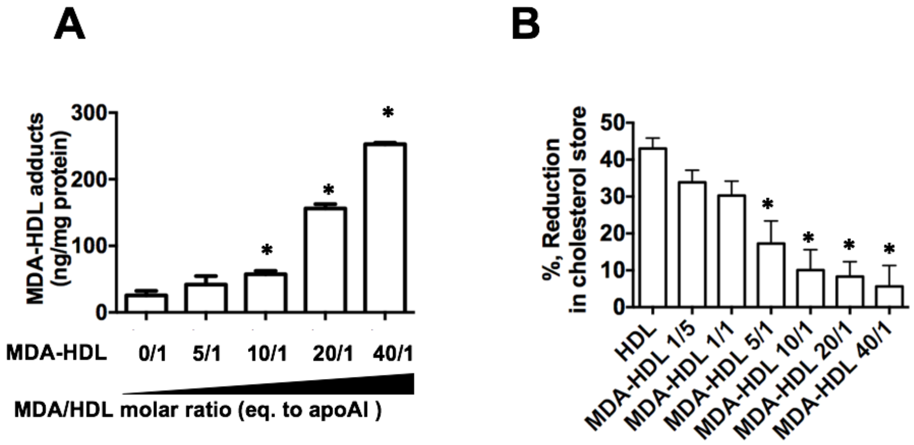


Supplemental Figure 12. Modification of HDL with increasing concentrations of MDA impaired cholesterol efflux in a dose dependent manner. (A) The HDL was modified with MDA, and the MDA adduct levels were measured by ELISA. (B) *Apoe*$^{-/-}$ peritoneal macrophages were incubated with ac-LDL for 40h and then incubated for 24h with 50ug/mL of HDL or MDA-HDL. The net cholesterol efflux capacity was measured as described in methods (One-way ANOVA with Bonferroni's Post-test, * indicates $p<0.05$).
